# Distinct modes of redox-mediated DegP activation govern its role as a general protease

**DOI:** 10.1101/2025.11.03.686187

**Authors:** Deeptodeep Roy, Ardra Nandakumar, Rachna Chaba

## Abstract

DegP, a bifunctional chaperone-protease central to maintaining envelope proteostasis in Gram-negative bacteria, serves as a paradigm for understanding protein quality control mechanisms. Decades of research, mainly using synthetic peptides/non-native substrates that do not contain cysteines, have established the allosteric mode of DegP activation, where substrate binding remodels inactive DegP into its active conformations. In enteric bacteria, as DegP typically harbours a disulfide bond (DSB) in its protease domain, its redox-mediated activation has also been proposed. However, the cellular factors driving this activation mode and its physiological relevance have remained elusive. Here, we investigated this aspect in *Escherichia coli* primarily using long-chain fatty acids and alkaline pH as stressors. We show that, under these conditions, DegP considerably accumulates in its thiol form (DegP_red_), which represents its protease-active conformation. Besides removing damaged proteins, DegP_red_ also induces the Cpx envelope stress response in all tested conditions. Notably, two distinct mechanisms promote redox-mediated DegP activation in a temporal and stress-specific manner: a compromised DSB-forming machinery fails to oxidize DegP, and thiol-containing substrates allosterically convert DegP into DegP_red_. Because *in vitro*, the lack of DSB sensitizes DegP to lower substrate levels, we suggest that *in vivo* DegP_red_ is primed for a robust response to stresses. Finally, the diverse nature of the molecular players that activate Cpx in our tested stress conditions leads us to propose that the DSB of DegP, by acting as a redox sensor, enables DegP to integrate a wide range of inducing cues and thus work as a general protease.

**Significance Statement:** The proteolytic activity of DegP is highly implicated in bacterial survival and virulence. Notably, DegP from the *Enterobacteriaceae* family of Gram-negative bacteria typically harbours a disulfide bond (DSB) in its protease domain. Because pathogens face redox stress during infection and antibiotic exposure, understanding how redox perturbants modulate DegP function is necessary for exploiting it as a target for therapeutic intervention strategies. Here, using *E. coli* as the model bacterium, we show that the DSB of DegP acts as a redox sensor under various inducing cues, thereby positioning DegP as a versatile, broad-spectrum protease. The present study opens avenues for investigating whether a similar redox-mediated control of DegP activation influences the pathogenesis of enteric bacteria.

## Introduction

Protein misfolding threatens cellular fitness in all forms of life. In bacteria, protein misfolding is of utmost concern in the envelope, an extracytoplasmic compartment that serves as the first line of defense against environmental insults and is the site for myriad critical processes (1). In the Gram-negative bacterium *E. coli*, the peripheral inner membrane protein DegP, a classical member of the HTRA (High temperature requirement protease A) family of proteins and a regulon member of the Cpx and σ^E^ envelope stress response (ESR) pathways, is a major protein quality control factor that functions both as a chaperone and a protease to maintain envelope integrity (2-6).

Under basal conditions, DegP is an inactive hexamer (DegP_6_) of two stacked trimers. Structurally, each DegP monomer constitutes an N-terminal protease domain carrying the catalytic serine (S210) and several loops of regulatory function, and two PDZ domains. In DegP_6_, the LA loop from one monomer interacts with the LA loop and the L1-L2 loops from another monomer of an opposing trimer, distorting the DegP catalytic site and keeping it inactive. Extensive studies on DegP have proposed a structure-driven functional model for the two widely prevalent modes of DegP activation. Briefly, upon the binding of misfolded substrates (allosteric model) and/or exposure to high temperatures, the inhibitory LA loop interactions are disturbed, resulting in structural remodeling that sets up the functional active site and transitions DegP_6_ to large active cage-like oligomers (DegP_12/24_) via trimeric intermediate (DegP_3_) (7-13) (Fig 1A and B). Notably, the *in vitro* studies leading to these activation models mainly used synthetic peptides/non-native substrates that do not contain cysteines.

**Figure 1.**
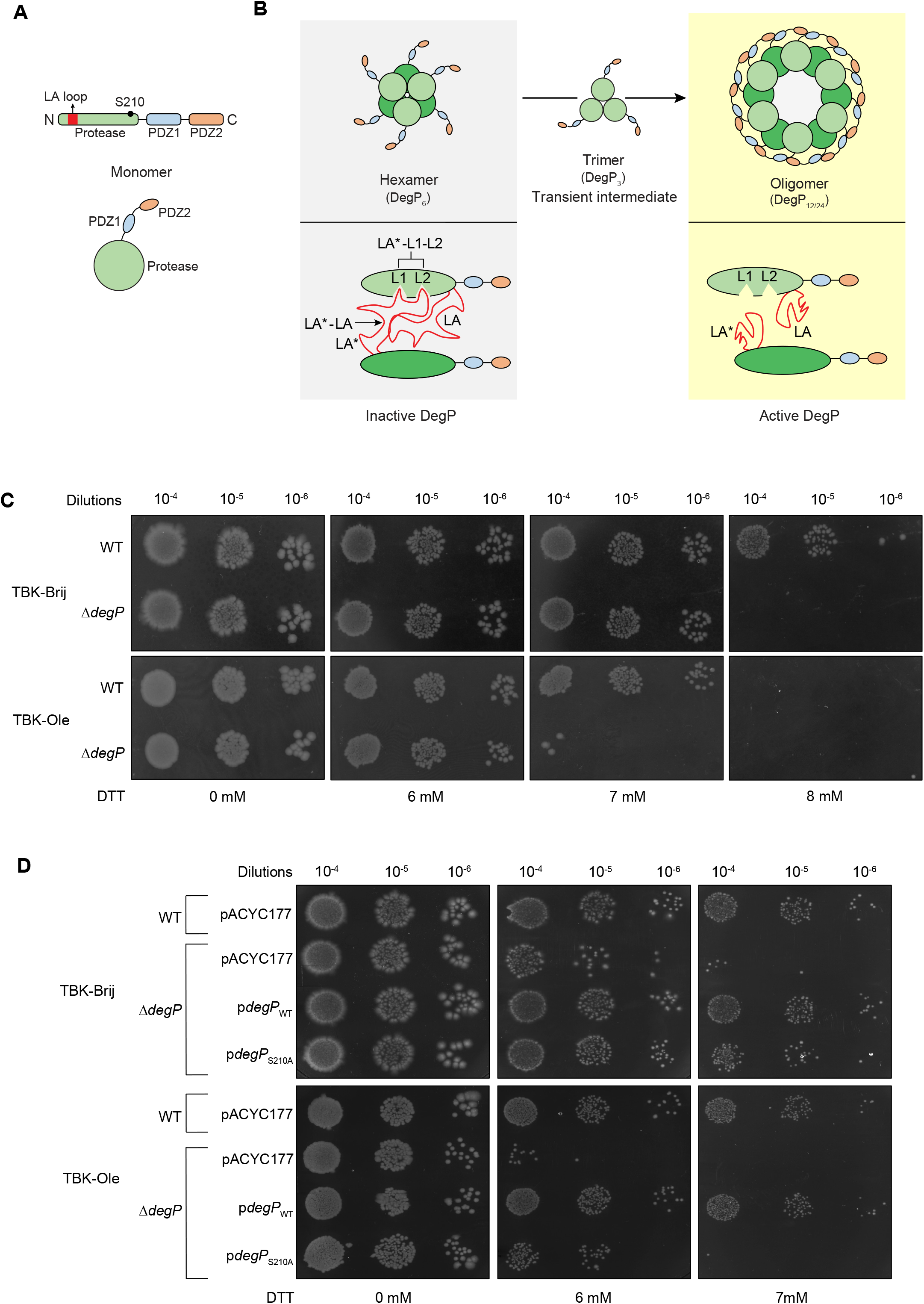
The proteolytic activity of DegP increases the fitness of E. coli exposed to thiol stress. (A) Schematic showing the linear (upper panel) and conformational (lower panel) forms of DegP monomer. Different colors indicate the various domains of DegP (13). (B) Schematic showing the structure-driven functional model of DegP activation. The upper panel shows the conformational states of DegP, and the lower panel depicts the interactions between the regulatory loops of different monomers. The protease domain of monomers from the two trimers are indicated in different shades of green. In DegP_6_, the LA loop from one monomer (LA*) interacts with the LA loop (LA*-LA) and the L1-L2 loops (LA*-L1-L2) from another monomer of an opposing trimer. DegP_6_ transitions to the active cage-like DegP_12/24_ via the DegP_3_ intermediate. In the active form, the LA*-LA interactions and LA*-L1-L2 triad are disrupted (13). (C) The Δ*degP* strain grown in oleate is hypersensitive to DTT. WT and Δ*degP* strains were spotted on TBK-Brij and TBK-Ole media with or without the indicated concentrations of DTT. The experiment was performed three times. A representative dataset is shown. (D) Oleate-grown cells expressing the protease-inactive form of DegP (DegP_S210A_) are hypersensitive to DTT. WT and Δ*degP* strains transformed with either pACYC177, pACYC177 carrying *degP* (p*degP*_WT_), or pACYC177 carrying *degP*_S210A_ (p*degP*_S210A_) were spotted on TBK-Brij and TBK-Ole media supplemented with or without the indicated concentrations of DTT. The experiment was performed twice. A representative dataset is shown.

An alternative mode of DegP activation, suggested by only a couple of studies, relies on its redox state. DsbA, a periplasmic oxidoreductase, which catalyzes disulfide bond (DSB) formation in secreted envelope proteins, introduces the only DSB in DegP, present in its LA loop. In contrast to wild-type (WT) cells, where DegP functions as a protease at high temperatures, in a Δ*dsbA* strain, it functions as a protease even at low temperatures, and compared to the WT protein, a cysteine-less mutant of DegP (ΔCys DegP) is a more efficient protease (14-16). Despite several decades of research that have established DegP as a paradigm in mitigating envelope problems, cellular factors promoting this redox-mediated activation of DegP and its physiological relevance remain elusive.

In *E. coli*, under aerobic conditions, the DSB-forming machinery is re-oxidized by ubiquinone, a mobile electron carrier in the electron transport chain (ETC) (17). Because ubiquinone also shuttles electrons derived from carbon metabolism, we previously demonstrated that an increased electron flow in the ETC during long-chain fatty acid (LCFA) metabolism limits ubiquinone availability for DSB formation, bringing DsbA into its thiol state when cells enter the stationary phase. Importantly, the Cpx ESR is activated in the stationary phase, which likely re-oxidizes DsbA and alleviates envelope stress (18-21). Because DegP is a DsbA substrate and a Cpx regulon member, LCFA metabolism represents an ideal condition for probing the factors that drive redox-mediated activation of DegP and understanding its physiological significance. Further, although DegP contributes to envelope integrity under various stress conditions (6, 22-25), the redox status of DegP and how it affects its function under these conditions have not been investigated.

Using LCFA metabolism and alkaline pH as inducing cues, here we conclusively establish that the thiol form of DegP (DegP_red_) represents its protease-active conformation. In LCFA-utilizing cells, whereas ubiquinone limitation is the reason for the accumulation of DegP_red_ till cells enter the stationary phase, in LCFA-grown stationary phase cells and in alkaline pH, DegP_red_ accumulates in a ubiquinone-independent manner, indicating that distinct factors drive redox-mediated activation of DegP. Further, by employing three additional redox perturbants i.e., *dsbA* deletion, DTT treatment, and overexpression of the outer membrane lipoprotein NlpE, our data suggest that besides clearing out aberrant proteins, DegP_red_ also contributes to Cpx activation. Because Cpx upregulates DegP, the *de novo* synthesized DegP is likely allosterically converted into DegP_red_ by misfolded proteins that accumulate under the above stressors. Finally, although DegP_red_ accumulation under these diverse environmental cues converges on Cpx activation, the distinct molecular players/signals activating Cpx lead us to speculate that the substrates modulating DegP’s redox state and subsequently targeted by the protease are condition-specific. This supports the role of DegP as a general protease responsive to a range of stress cues.

## Results

### Proteolytic activity of DegP is increasingly required in LCFA-grown cells to tackle redox stress

We previously showed that *E. coli* utilizing LCFAs accumulates DsbA and its substrate, DegP, considerably in the thiol state. Briefly, we monitored the redox state of DsbA and DegP from the exponential phase to entry into the stationary phase. Whereas DsbA was observed in its thiol form only in TBK-Ole-grown cells [cells grown in buffered tryptone broth (TBK) supplemented with oleate (C18 LCFA)] during entry into the stationary phase, DegP_red_ was observed to increase gradually from the exponential phase to entry into the stationary phase, both in TBK-Brij (TBK supplemented with Brij-58, a detergent used for solubilizing oleate; basal medium) and TBK-Ole-grown cells. However, at every time point, DegP_red_ was remarkably higher in TBK-Ole-grown cells (18). Because DegP, lacking its DSB, either in a Δ*dsbA* strain or in ΔCys DegP, functions as an efficient protease (14, 15), the presence of a higher amount of DegP_red_ in TBK-Ole compared to TBK-Brij led us to ask whether DegP plays a critical role during LCFA metabolism.

We previously showed that because LCFA metabolism generates redox stress, *E. coli* grown in TBK-Ole is more sensitive to the thiol agent, DTT, compared to cells grown in TBK-Brij (18). Here, we reasoned that if the higher levels of DegP_red_ in LCFA-grown cells combat redox stress, then a Δ*degP* strain cultured in TBK-Ole should be hypersensitive to DTT compared to both the WT strain grown in TBK-Ole and also the WT and Δ*degP* strains grown in TBK-Brij. Indeed, whereas in TBK-Brij, both WT and Δ*degP* grew similarly up to 7 mM DTT and Δ*degP* showed a growth defect at 8 mM DTT, in TBK-Ole, compared to WT, Δ*degP* exhibited a significant growth defect at 7 mM DTT while both WT and Δ*degP* could not grow at 8 mM DTT (Fig 1C). Collectively, these observations suggest that DegP provides cellular fitness under reducing conditions, and its requirement increases with an increase in redox stress, being higher in LCFA-grown cells, a redox stress-inducing condition.

Because DegP can function both as a chaperone and a protease (5), we next addressed whether the chaperone or the protease activity of DegP contributes to cellular fitness during redox stress. For this, we used a protease-inactive DegP_S210A_ mutant; DegP_S210A_ retains its chaperone/chaperone-like activity with no observable proteolytic activity (5). We checked the thiol sensitivity of WT and Δ*degP* strains expressing either WT DegP or DegP_S210A_ from pACYC177 in both TBK-Brij and TBK-Ole. The expression of WT DegP from pACYC177 fully rescued the growth defect of Δ*degP* in TBK-Brij and TBK-Ole supplemented with either 6 mM or 7 mM DTT; however, a distinct pattern was observed upon expression of DegP_S210A_. In TBK-Brij, DegP_S210A_ completely rescued the growth defect at 6 mM DTT but only allowed partial rescue at 7 mM DTT. In TBK-Ole, whereas DegP_S210A_ enabled partial rescue at 6 mM DTT, it could not rescue the growth defect at 7 mM DTT (Fig 1D). WT DegP and DegP_S210A_ expressed to a similar level from pACYC177 (Fig S1), validating that their differential ability to counter redox stress is due to their distinct functionality and not due to a difference in protein levels. These data suggest that while the chaperone activity retained in DegP_S210A_ can sustain cells under low redox stress, the proteolytic activity becomes critical under high redox stress, with its requirement being higher in LCFA-grown cells, a redox-perturbing condition. Taken together, these data establish the critical role of DegP in providing fitness during LCFA metabolism.

### Proteolytic activity of DegP participates in Cpx activation under redox-perturbing conditions

DegP is widely known to relieve different kinds of stresses by degrading a broad range of stress-induced misfolded substrates (6, 22, 24, 26). However, in alkaline pH, DegP also counteracts stress by participating in Cpx activation (23). Because Cpx is induced in LCFA-grown cells (18), here we investigated whether DegP has any role in its activation. Using a *cpxP*-*lacZ* chromosomal transcriptional reporter where *lacZ* was fused downstream of the promoter of *cpxP* [*cpxP* is a *bona fide* Cpx regulon member (27)], we observed that, whereas in the WT strain, Cpx was induced ∼3.5-fold in TBK-Ole in the stationary phase (time point T5, Fig 2A), in the Δ*degP* strain, Cpx response was reduced to ∼2-fold, i.e., there was ∼40% decrease in Cpx induction in the absence of DegP (Fig 2B). These data prompted us to examine whether DegP also activates Cpx in other redox-perturbing conditions. For this, we first determined the effect of *degP* deletion on Cpx activation in a Δ*dsbA* strain and DTT-treated cells. The ∼3-fold increase in Cpx activation observed in a Δ*dsbA* strain decreased to ∼2-fold upon *degP* deletion (Fig 2C), and the ∼2-fold increase in Cpx response observed in WT cells upon DTT treatment reduced to ∼1.5-fold in the Δ*degP* strain (Figs S2 and 2D). We further probed the role of DegP in activating Cpx in cells overexpressing NlpE, a well-known Cpx inducer and a sensor of oxidative protein folding defects (28-30). In NlpE overexpressing cells, Cpx activity exhibited a marginal, yet significant decrease upon *degP* deletion, i.e., ∼12-fold in the WT strain vs. ∼10-fold in the Δ*degP* strain (Fig 2E). Notably, similar to LCFA-grown cells, the Δ*dsbA* strain, and DTT-treated cells (15, 16, 18, 31), NlpE overexpression remarkably brought DegP into its thiol form (Fig 2F).

**Figure 2.**
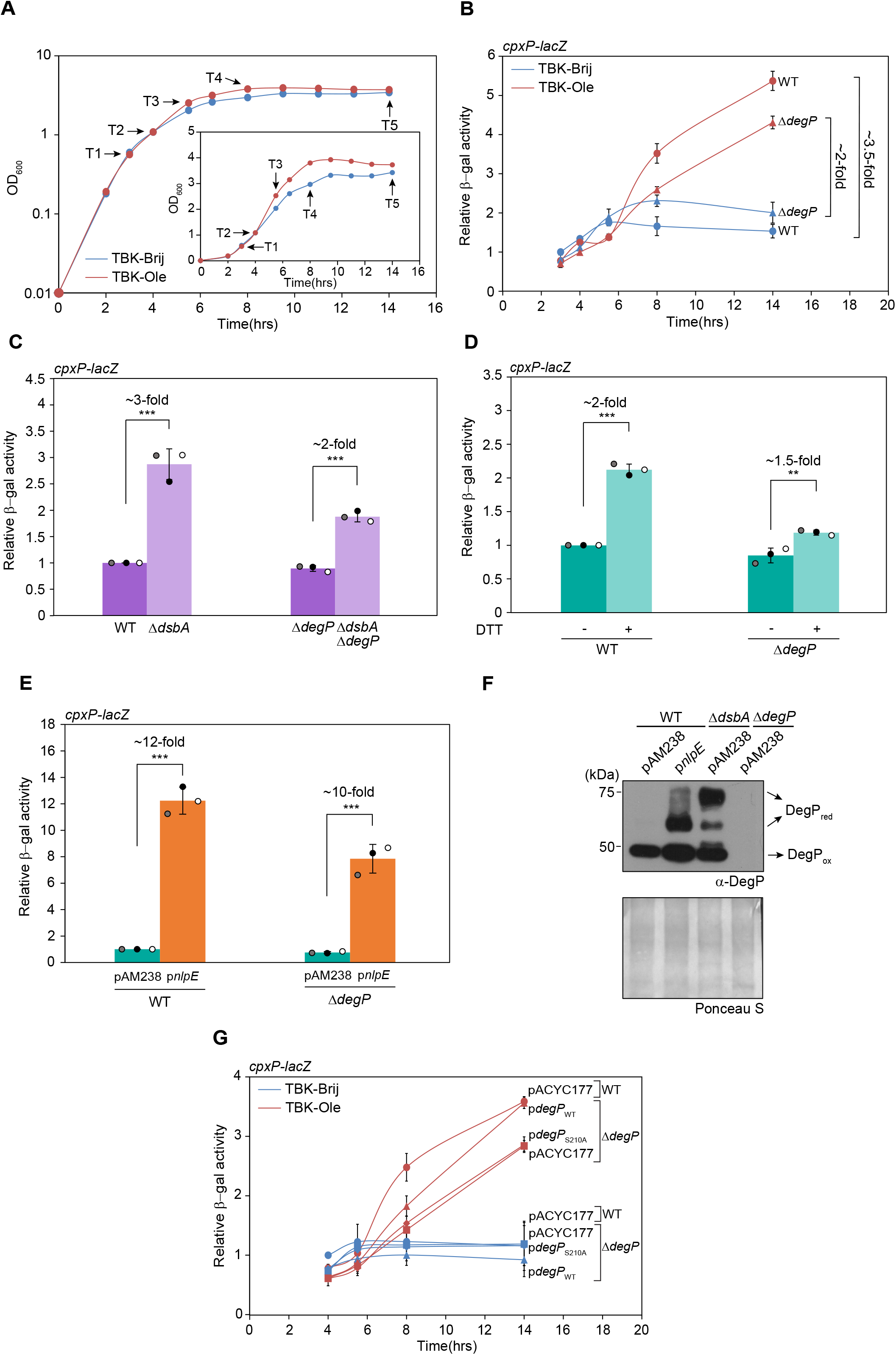
E. coli facing redox stress requires the proteolytic activity of DegP for Cpx activation. (A) Growth curve of WT strain in basal and oleate-supplemented media. OD_600_ of the cultures was measured, and the growth curve was plotted on a semi-logarithmic scale. T1, T2, T3, T4, and T5 indicate time points where cultures were harvested for various assays. The experiment was done three times. A representative dataset is shown. *Inset:* The growth curve was also plotted on a linear scale. (B) Cpx induction significantly decreases in the Δ*degP* strain grown in oleate. Strains carrying *cpxP-lacZ* transcriptional reporter were grown either in TBK-Brij or TBK-Ole. Cultures were harvested from different phases of growth corresponding to the time points as indicated in Fig 2A, and β-gal activity was measured. Data were normalized to the β-gal activity of WT in TBK-Brij at time point T1. Data represent the average (±S.D.) of three independent experiments. The average β-gal activity (in Miller units) of WT *cpxP-lacZ* in TBK-Brij at time point T1 was 36 (±5). (C) DegP contributes to Cpx activation in a Δ*dsbA* strain. Strains carrying *cpxP-lacZ* transcriptional reporter were grown in LB. Cultures were harvested from the exponential phase, and β-gal activity was measured. Data were normalized to the β-gal activity of WT. Data represent the average (±S.D.) of three independent experiments. The average β-gal activity (in Miller units) of WT *cpxP-lacZ* was 6 (±1). (D) DegP contributes to Cpx activation in DTT-treated cells. Strains carrying *cpxP-lacZ* transcriptional reporter were grown in TBK supplemented with or without 1.8 mM DTT. Cultures were harvested after three hours of growth (as indicated in Fig S2), and β-gal activity was measured. Data were normalized to the β-gal activity of WT grown without DTT. Data represent the average (±S.D.) of three independent experiments. The average β-gal activity (in Miller units) of WT *cpxP-lacZ* grown without DTT was 51 (±8). (E) DegP participates in Cpx activation upon NlpE overexpression. Strains carrying *cpxP-lacZ* transcriptional reporter were transformed with either pAM238 or pAM238 carrying *nlpE* (p*nlpE*) and were grown in TBK medium supplemented with 0.1 mM IPTG. Cultures were harvested in the exponential phase, and β-gal activity was measured. Data were normalized to the β-gal activity of WT carrying pAM238 and represent the average (±S.D.) of three independent experiments. The average β-gal activity (in Miller units) of WT *cpxP-lacZ* carrying pAM238 was 33 (±6). (F) DegP significantly accumulates in its thiol state upon NlpE overexpression. Strains were transformed with either pAM238 or p*nlpE* and were grown in TBK medium supplemented with 0.1 mM IPTG. Cultures were harvested in the exponential phase. Proteins were denatured and precipitated using trichloroacetic acid, followed by treatment with MAL-PEG, a maleimide derivative that irreversibly alkylates free thiols, adding ∼5 kDa per thiol group (54). Oxidized and reduced forms of DegP were identified by running the samples on non-reducing SDS-PAGE gels, followed by Western blotting using an anti-DegP antibody. Δ*dsbA* and Δ*degP* strains transformed with pAM238 served as controls. DegP_ox_ and DegP_red_ indicate oxidized and reduced forms of DegP, respectively. DegP is oxidized by DsbA (15, 16), hence DegP accumulates as DegP_red_ in the Δ*dsbA* strain. Ponceau S-stained counterpart of the Western blot served as a loading control. The blot shown is a representative of three independent replicates. (G) In oleate, the protease-inactive mutant of DegP exhibits Cpx induction similar to the Δ*degP* strain. Strains carrying *cpxP-lacZ* transcriptional reporter transformed with either pACYC177, p*degP*_WT_, or p*degP*_S210A_ were grown either in TBK-Brij or TBK-Ole. Cultures were harvested at different phases of growth corresponding to the time points as indicated in Fig 2A, and β-gal activity was measured. Data were normalized to the β-gal activity of WT carrying pACYC177 in TBK-Brij at time point T2. Data represent the average (±S.D.) of three independent experiments. The average β-gal activity (in Miller units) of WT carrying pACYC177 in TBK-Brij at time point T2 was 53 (±9). For panels C, D and E, the p-values were calculated using the unpaired two-tailed Student’s t-test (***, P<0.001; **, P<0.01).

Both the presence of DegP as DegP_red_ and the involvement of DegP in Cpx activation under all tested redox-perturbing conditions indicated that DegP_red_ activates Cpx. Further, the increased accumulation of DegP_red_ and the increased requirement of the proteolytic activity of DegP with increasing redox stress suggested a link between DegP_red_ and its protease activity (18) (Fig 1D). Collectively, these correlations indicated that the proteolytic activity of DegP is involved in Cpx activation under redox stress. We investigated this aspect, taking LCFA metabolism as a representative condition. In TBK-Ole, whereas Δ*degP* expressing WT DegP from pACYC177 showed Cpx response similar to that of the WT strain, the Cpx response of Δ*degP* expressing DegP_S210A_ from pACYC177 was similar to that of the Δ*degP* strain. These data establish that the proteolytic activity of DegP is involved in Cpx induction (Fig 2G), thereby emphasizing that besides clearing out damaged proteins, DegP provides cellular fitness in redox-perturbing conditions by inducing the Cpx response.

### DegP-mediated Cpx activation during LCFA metabolism is independent of CpxP degradation

The Cpx ESR comprises an inner membrane sensor histidine kinase CpxA, a cytoplasmic response regulator CpxR, and a periplasmic auxiliary component CpxP that binds CpxA and acts as a negative regulator. During envelope stress, CpxA autophosphorylates and transphosphorylates CpxR, which in turn directs the transcription of its regulon members involved in mitigating stress (32, 33). In alkaline pH, DegP degrades CpxP, and Cpx activation is partially DegP-dependent (23), indicating that DegP-mediated proteolysis of CpxP relieves its inhibition of the Cpx response. We, therefore, asked if DegP degrades CpxP to activate Cpx in LCFA-grown cells as well. Because CpxP is a negative regulator of the Cpx pathway, in TBK-Brij, we expected a higher Cpx activity in a Δ*cpxP* strain. Further, we expected that in TBK-Ole, Cpx activity should either be similar in WT and Δ*cpxP* strains (if DegP degrades CpxP completely) or the fold increase in Δ*cpxP* vs. WT in TBK-Ole should be lower than the fold increase in TBK-Brij (if DegP degrades CpxP partially). Surprisingly, the fold increase in Cpx activation in Δ*cpxP* vs. WT was similar (1.7-fold) in both the media conditions (Fig 3A), providing the first indication that CpxP is not involved in Cpx activation during LCFA metabolism. Notably, corroborating with the remarkably higher Cpx induction in Δ*cpxP* strain in TBK-Ole, deletion of *cpxP* provided fitness advantage to LCFA-grown cells exposed to thiol stress; in 8 mM DTT, the Δ*cpxP* strain grew better than WT in TBK-Ole and both WT and Δ*cpxP* in TBK-Brij (Fig S3). Next, we compared Cpx induction in Δ*cpxP* and Δ*cpxP*Δ*degP* strains. We expected that if DegP-mediated Cpx activation proceeds through CpxP proteolysis, then in TBK-Ole, the extent of Cpx activation should be similar in Δ*cpxP* and Δ*cpxP*Δ*degP* strains. However, Cpx activation in Δ*cpxP*Δ*degP* was significantly lower than Δ*cpxP* (Figs 3B and S4), reiterating that in LCFA-grown cells, Cpx activation by DegP is independent of CpxP. As a final test, we directly determined the levels of CpxP in TBK-Brij and TBK-Ole-grown cells. Because in LCFA-grown cells, degradation of chromosomal CpxP by DegP, if any, would decrease its levels, whereas Cpx activation would increase its levels, to circumvent this issue, we expressed CpxP from a heterologous promoter from a plasmid. We observed comparable CpxP levels in TBK-Brij and TBK-Ole-grown cells (Fig 3C). Taken together, our data convincingly establish that, unlike alkaline pH, in LCFA-grown cells, DegP-mediated activation of Cpx is CpxP-independent.

**Figure 3.**
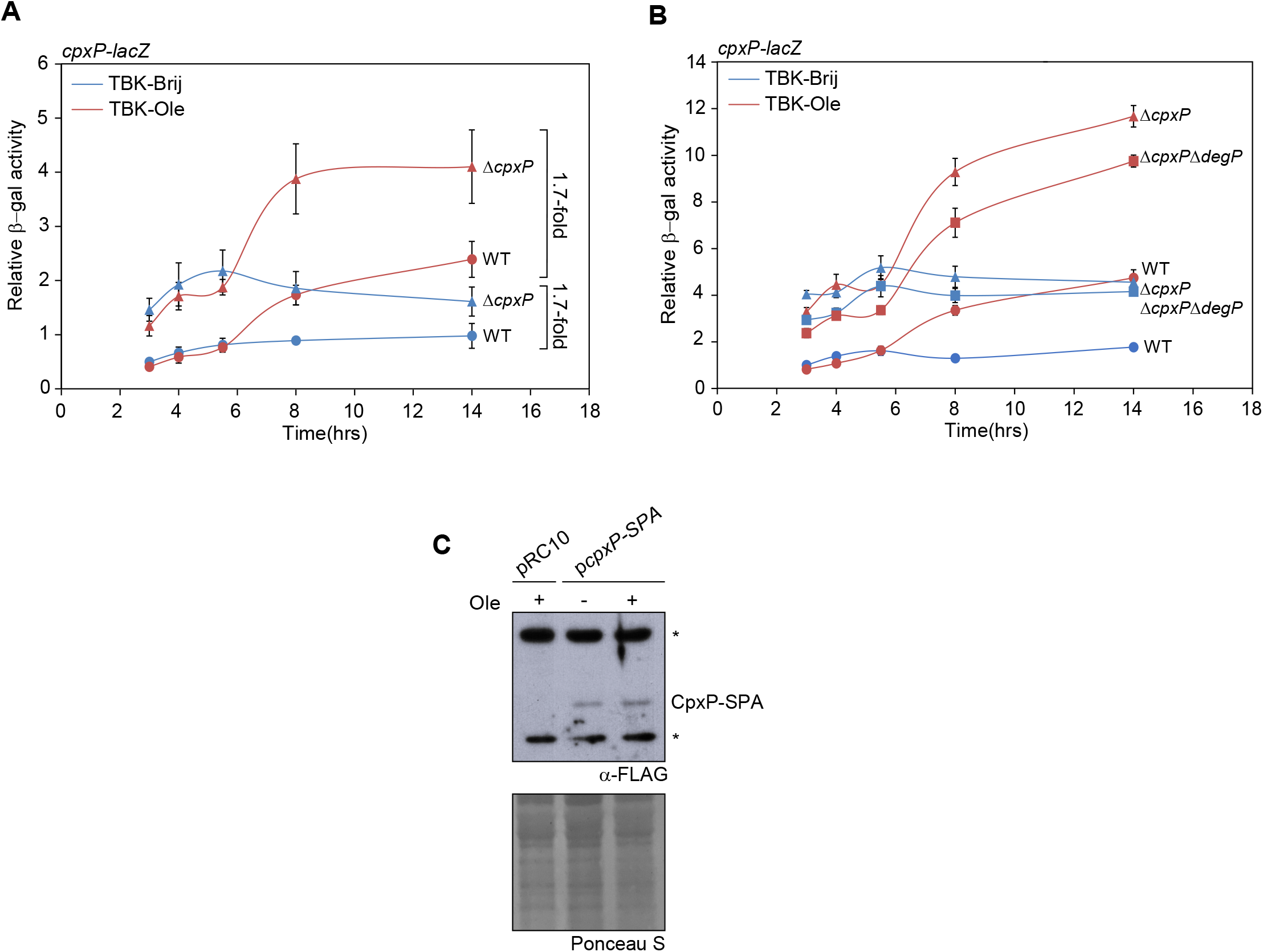
DegP activates Cpx in LCFA-grown cells independent of CpxP degradation. (A) In oleate-grown cells, CpxP is not involved in Cpx activation. Strains carrying *cpxP-lacZ* transcriptional reporter were grown either in TBK-Brij or TBK-Ole. Cultures were harvested at different phases of growth corresponding to the time points as indicated in Fig 2A, and β-gal activity was measured. Data were normalized to the β-gal activity of WT grown in TBK-Brij at time point T1 and represent the average (±S.D.) of three independent experiments. The average β-gal activity (in Miller units) of WT *cpxP-lacZ* in TBK-Brij at time point T1 was 39 (±1). (B) In oleate-grown cells, DegP-mediated activation of Cpx is CpxP-independent. Strains carrying *cpxP-lacZ* transcriptional reporter were grown either in TBK-Brij or TBK-Ole. Cultures were harvested in triplicate at different phases of growth corresponding to the time points as indicated in Fig 2A, and β-gal activity was measured. Data were normalized to the β-gal activity of WT grown in TBK-Brij at time point T1 and represent the average (±S.D.) of three technical replicates from one experiment. The average β-gal activity (in Miller units) of WT *cpxP-lacZ* in TBK-Brij at time point T1 was 50 (±4). Two independent experiments were performed; although the fold change varied across biological replicates, the trend observed was the same. Data from another independent experiment is shown in Fig S4. (C) CpxP levels are similar in cells grown in basal and oleate-supplemented media. The WT strain transformed with either pRC10 or pRC10 carrying *cpxP-SPA* (p*cpxP-SPA*) was grown either in TBK-Brij or TBK-Ole. Cultures were induced with 0.4 mM IPTG at time point T3, followed by harvesting at time point T4 (time points as indicated in Fig 2A). CpxP-SPA was probed using an anti-FLAG antibody. The band corresponding to CpxP-SPA (∼26 kDa) is shown. Ponceau S-stained counterpart of the Western blot served as a loading control. * indicates non-specific bands detected by anti-FLAG antibody, which serve as additional loading controls. The blot shown is a representative of two independent replicates.

### Ubiquinone limitation drives redox-mediated activation of DegP in LCFA-grown cells

Because DegP, as a protease, is involved in Cpx activation in LCFA-grown cells (Fig 2G), here we used Cpx as a readout to investigate the reason for its redox-mediated activation in this metabolic condition. In LCFA-utilizing cells, the exogenous supplementation of ubiquinone-8 (the natural form of ubiquinone in *E. coli*) lowers the Cpx response by ∼40%, suggesting that this fraction of Cpx activation is due to ubiquinone limitation (18) (Fig 4A). Further, cells lacking either DegP or expressing DegP_S210A_ also display a similar decrease in Cpx response (Fig 2B and G). Together, these findings suggest that ubiquinone limitation brings DegP into its protease-active form, which participates in Cpx activation. Consistent with this model, we found that Cpx induction is similar in a Δ*degP* strain, regardless of ubiquinone supplementation (Fig 4A).

**Figure 4.**
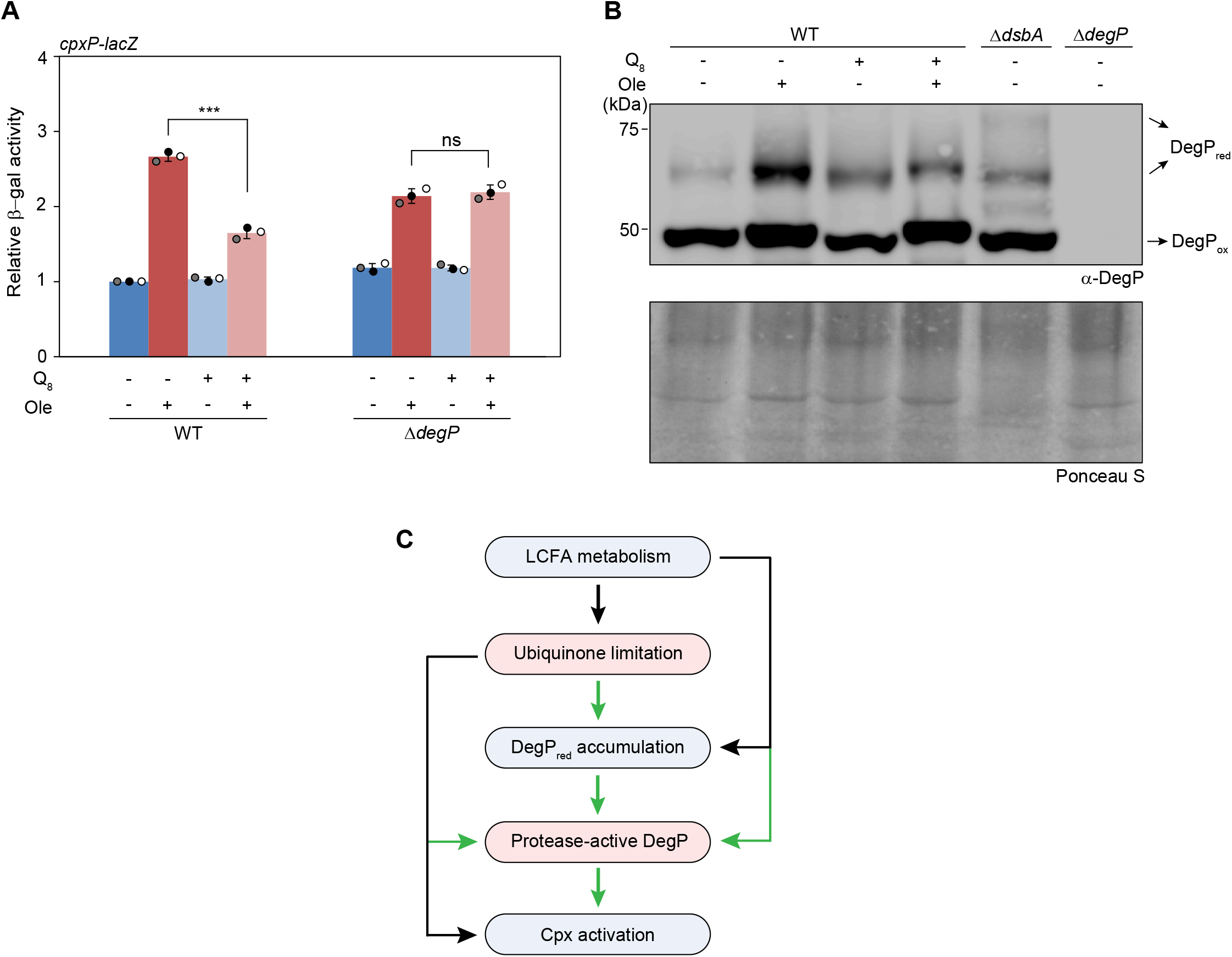
Ubiquinone limitation activates DegP in LCFA-grown cells. (A) In oleate-grown cells, ubiquinone limitation activates the Cpx pathway via DegP. Strains carrying *cpxP-lacZ* transcriptional reporter were grown either in TBK-Brij (-Ole) or TBK-Ole (+Ole). The media contained either 20 µM ubiquinone-8 (Q_8_+) or 0.1% ethanol (Q_8_-), the solvent used for solubilizing ubiquinone-8, as indicated. Cultures were harvested in the stationary phase (time point T5, as indicated in Fig S5) and β-gal activity was measured. Data were normalized to the β-gal activity of WT grown in TBK-Brij containing 0.1% ethanol and represent the average (±S.D.) of three independent experiments. The average β-gal activity (in Miller units) of WT *cpxP-lacZ* in TBK-Brij supplemented with 0.1% ethanol was 88 (±16). The p-values were calculated using the unpaired two-tailed Student’s t-test (***, P<0.001; ns, P>0.05). (B) Ubiquinone limitation brings DegP into its thiol form in oleate-grown cells. Strains were grown either in TBK-Brij (-Ole) or TBK-Ole (+Ole). The media contained either 20 µM ubiquinone-8 (Q_8_+) or 0.1% ethanol (Q_8_-), as indicated. Cells were harvested in the stationary phase (time point T5, as indicated in Fig S5) and processed as mentioned in the legend to Fig 2F, followed by probing with an anti-DegP antibody. Δ*dsbA* and Δ*degP* strains cultured in TBK-Brij containing 0.1% ethanol served as controls. DegP_ox_ and DegP_red_ indicate oxidized and reduced forms of DegP, respectively. Ponceau S-stained counterpart of the Western blot served as a loading control. The blot shown is a representative of three independent replicates. (C) In LCFA-grown cells, ubiquinone limitation results in the accumulation of DegP as DegP_red_, which functions as a protease to activate Cpx. The black lines indicate findings from our previous work (18). The green lines indicate observations from the current study.

Next, we directly examined the redox state of DegP. In LCFA-grown cells, the accumulation of DsbA in its thiol state during entry into the stationary phase is due to ubiquinone limitation (18). Because DegP is a DsbA substrate (15, 16), here we examined whether ubiquinone limitation is also the reason for the considerable accumulation of DegP_red_ in LCFA-grown cells. For this, we determined the redox state of DegP in WT cells grown in TBK-Brij and TBK-Ole, in the presence or absence of ubiquinone. Considering that the addition of ubiquinone attenuated the DegP-mediated Cpx response in the stationary phase (Fig 4A), we determined the redox state of DegP in this growth phase. Whereas in the absence of ubiquinone supplementation, there was a considerable accumulation of DegP_red_ in cells grown in TBK-Ole compared to TBK-Brij, upon ubiquinone supplementation, accumulation of DegP_red_ decreased significantly in TBK-Ole (Fig 4B). The considerable accumulation of DegP_red_ in TBK-Ole compared to TBK-Brij is not due to a difference in growth in these media; the growth profile of WT in TBK-Ole with or without ubiquinone supplementation was the same, yet DegP was considerably present as DegP_red_ without ubiquinone supplementation compared to upon ubiquinone supplementation (compare Figs 4B and S5). Collectively, our data clearly show that ubiquinone limitation leads to a considerable accumulation of DegP_red_ during LCFA metabolism.

Because ubiquinone limitation brings DegP into its protease-active form (Fig 4A) and also its thiol form (Fig 4B), these data validate DegP_red_ as the protease form and convincingly establish ubiquinone limitation as the reason for the redox-mediated activation of DegP in LCFA-grown cells (Fig 4C).

### Accumulation of DegP in its thiol form under alkaline stress is ubiquinone-independent

Previous observations that Cpx activation in alkaline pH is partly DegP-dependent and CpxP is detected in a Δ*degP* strain but not in a WT strain collectively put forth a model that DegP degrades CpxP to activate Cpx (23). However, whether the proteolytic activity of DegP is indeed involved in Cpx activation was not directly investigated. Because in LCFA-metabolizing cells, DegP_red_ corresponds to its protease-active form, which activates Cpx (Fig 4), here we investigated whether, under alkaline stress as well, DegP exists as DegP_red_, which participates in Cpx activation. For this, we first determined Cpx induction in WT and Δ*degP* strains in acidic (pH 5.8), neutral (pH 6.8), and alkaline (pH 8.2) conditions during the exponential phase (Fig S6). Consistent with earlier reports, we found that DegP activates Cpx under alkaline pH; the Cpx response gradually increased with increasing pH and showed ∼30% decrease upon *degP* deletion only at pH 8.2 (Fig 5A). Importantly, in this condition, whereas WT DegP expressed from pACYC177 restored Cpx induction in the Δ*degP* strain to WT levels, the expression of DegP_S210A_ could not do so (Fig 5B). Further, assessing the redox state of DegP, we observed that the amount of DegP_red_ progressively increased with increasing pH, with a maximal accumulation at pH 8.2 (Fig 5C). Taken together, these data show that, similar to LCFA-metabolizing cells, under alkaline pH, where there is a considerable accumulation of DegP as DegP_red_, its proteolytic activity is involved in Cpx activation, reiterating that DegP_red_ represents its proteolytically active state.

**Figure 5.**
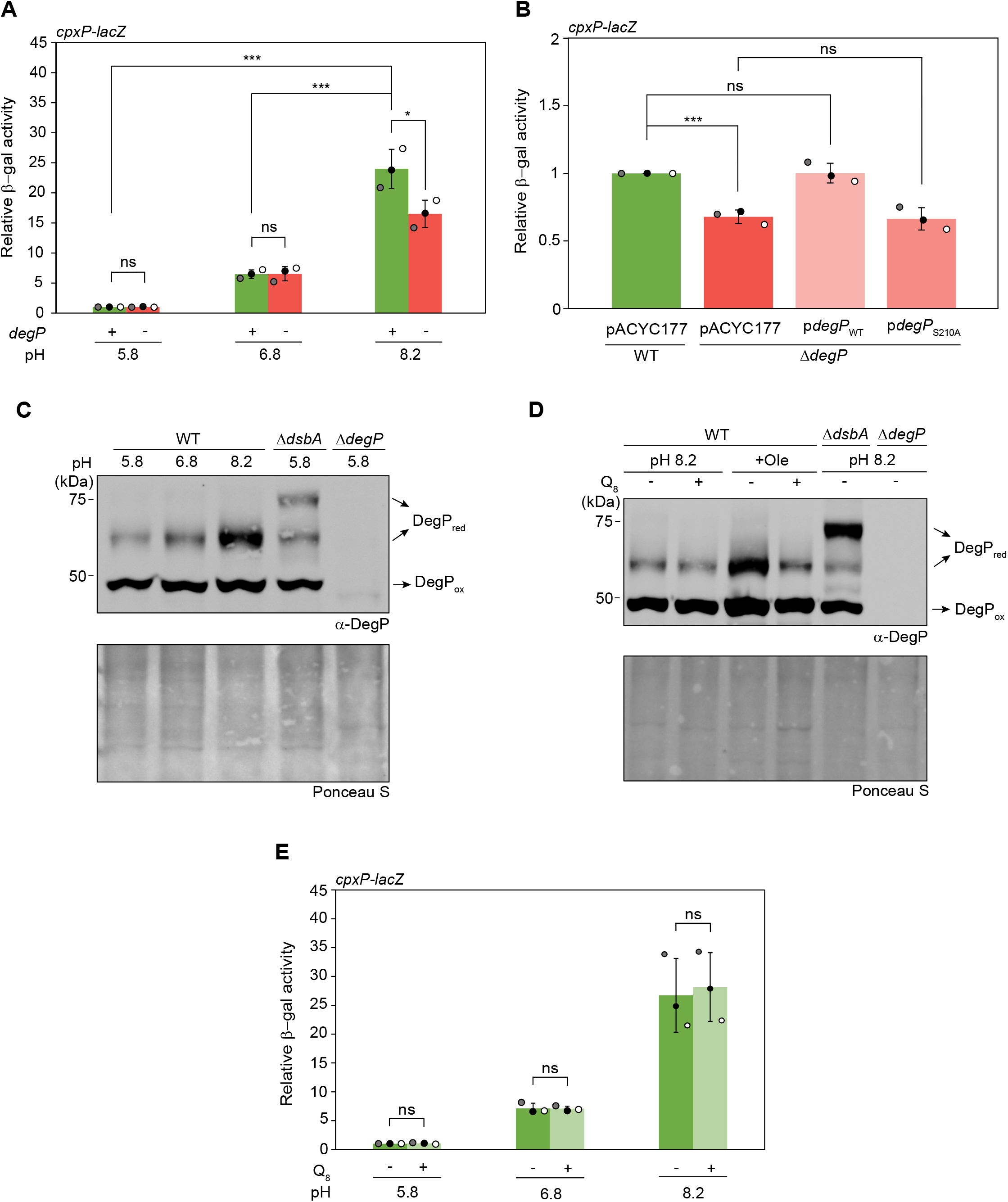
Activation of DegP under alkaline pH is ubiquinone-independent. (A) DegP is involved in Cpx activation in cells under alkaline pH stress. Strains carrying *cpxP-lacZ* transcriptional reporter were grown in LB of different pH, as indicated. Cultures were harvested in the exponential phase (as indicated in Fig S6) and β-gal activity was measured. Data were normalized to the β-gal activity of WT grown in LB of pH 5.8 and represent the average (±S.D.) of three independent experiments. The average β-gal activity (in Miller units) of WT *cpxP-lacZ* in LB of pH 5.8 was 5 (±1). (B) The proteolytic activity of DegP is required for Cpx activation in cells exposed to alkaline pH. Strains carrying *cpxP-lacZ* transcriptional reporter transformed with either pACYC177, p*degP*_WT_, or p*degP*_S210A_ were grown in LB of pH 8.2. Cultures were harvested in the exponential phase (as indicated in Fig S6), and β-gal activity was measured. Data were normalized to the β-gal activity of WT transformed with pACYC177 and represent the average (±S.D.) of three independent experiments. The average β-gal activity (in Miller units) of WT *cpxP-lacZ* transformed with pACYC177 was 133 (±8). (C) DegP_red_ maximally accumulates in alkaline pH. Strains were grown in LB of different pH, as indicated. Cells were harvested in the exponential phase (as indicated in Fig S6), and processed as mentioned in the legend to Fig 2F, followed by probing with an anti-DegP antibody. Δ*dsbA* and Δ*degP* strains cultured in pH 5.8 served as controls. DegP_ox_ and DegP_red_ indicate oxidized and reduced forms of DegP, respectively. Ponceau S-stained counterpart of the Western blot served as a loading control. The blot shown is a representative of two independent replicates. (D) Unlike LCFA-grown cells, DegP_red_ levels do not change in cells grown in alkaline pH upon ubiquinone supplementation. Strains were grown in LB of pH 8.2 or TBK-Ole (+Ole), as indicated. The media contained either 20 µM ubiquinone-8 (Q_8_+) or 0.1% ethanol (Q_8_-) as mentioned. Cultures grown in LB of pH 8.2 were harvested in the exponential phase (as indicated in Fig S6), and those grown in TBK-Ole were harvested in the stationary phase (time point T5, as indicated in Fig S5). Cells were processed as mentioned in the legend to Fig 2F, followed by probing with an anti-DegP antibody. Δ*dsbA* and Δ*degP* strains cultured in LB of pH 8.2 served as controls. DegP_ox_ and DegP_red_ indicate oxidized and reduced forms of DegP, respectively. Ponceau S-stained counterpart of the Western blot served as a loading control. The blot shown is a representative of two independent replicates. (E) Cpx induction in cells grown in alkaline pH is ubiquinone-independent. Strains carrying *cpxP-lacZ* transcriptional reporter were grown in LB of different pH, as indicated. The media contained either 20 µM ubiquinone-8 (Q_8_+) or 0.1% ethanol (Q_8_-) as indicated. Cultures were harvested in the exponential phase (as indicated in Fig S6), and β-gal activity was measured. Data were normalized to the β-gal activity of WT grown in LB of pH 5.8 without ubiquinone supplementation and represent the average (±S.D.) of three independent experiments. The average β-gal activity (in Miller units) of WT *cpxP-lacZ* in LB of pH 5.8 was 4 (±2). For panels A, B, and E, the p-values were calculated using the unpaired two-tailed Student’s t test (***, P<0.001; *, P<0.05; ns, P>0.05).

Considering that ubiquinone limitation brings DegP into its thiol form in LCFA-grown cells, we next investigated whether this factor also underlies DegP activation under alkaline conditions. However, neither DegP_red_ accumulation nor Cpx response at pH 8.2 changed upon ubiquinone supplementation (Figs 5D,E and S7). These data suggest that distinct factors govern redox-mediated activation of DegP under different stress conditions.

### Distinct factors contribute to the redox-mediated activation of DegP in a temporal manner in LCFA-grown cells

We previously showed that whereas DsbA entirely accumulates in its thiol form when LCFA-metabolizing cells enter the stationary phase, it is re-oxidized in the stationary phase. Further, our data showed that Cpx is activated in the stationary phase, which both upregulates ubiquinone and downregulates LCFA metabolism, suggesting that the Cpx pathway increases ubiquinone availability for re-oxidation of the DSB-forming machinery (18, 19). Because DegP is a DsbA substrate, the remarkable accumulation of DegP_red_ in the stationary phase, despite DsbA re-oxidation, was counterintuitive (Fig 4B). To address this discrepancy, we first performed a direct phase-wise comparison of the redox state of DegP in LCFA-metabolizing cells from the exponential to the stationary phase of growth (across time points T1 to T5, indicated in Fig 2A). Consistent with our previous results (18), from exponential phase to entry into the stationary phase (time points T1 to T3), although the DegP levels did not increase, DegP_red_ levels gradually increased with increasing cell density in both TBK-Brij and TBK-Ole-grown cells, with a higher accumulation in TBK-Ole at each time point (Fig 6A and B), indicating that the higher levels of DegP_red_ during LCFA metabolism are solely a consequence of a progressive increase in thiol stress, i.e., ubiquinone limitation.

**Figure 6.**
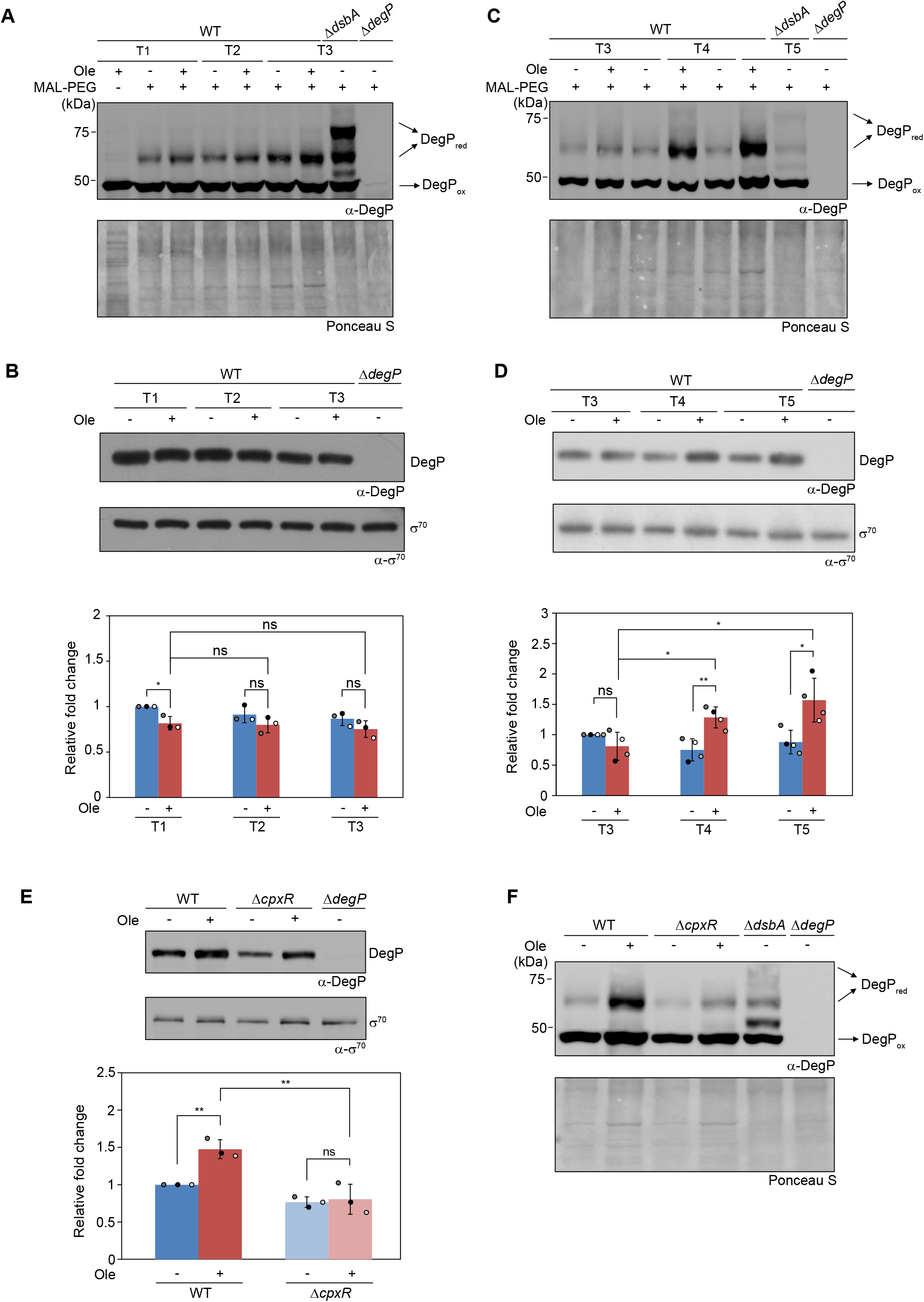
Both total DegP and DegP_red_ levels increase in stationary phase LCFA-grown cells in a Cpx-dependent manner. (A) The reduced form of DegP accumulates in oleate-grown cells till entry into the stationary phase. Strains were grown either in TBK-Brij (-Ole) or TBK-Ole (+Ole). Cells were harvested at different phases of growth corresponding to time points T1, T2, and T3 (as indicated in Fig 2A), and processed as mentioned in the legend to Fig 2F, followed by probing with an anti-DegP antibody. Δ*dsbA* and Δ*degP* strains cultured in TBK-Brij served as controls. DegP_ox_ and DegP_red_ indicate oxidized and reduced forms of DegP, respectively. The sample without MAL-PEG treatment only shows the DegP_ox_ form. Ponceau S-stained counterpart of the Western blot served as a loading control. Each blot is a representative of three independent replicates. (B) Total DegP levels in oleate-grown cells do not increase till entry into the stationary phase. Strains were grown either in TBK-Brij (-Ole) or TBK-Ole (+Ole). Cells were harvested at different phases of growth corresponding to time points T1, T2, and T3 (as indicated in Fig 2A). Lysates were prepared, samples were run on SDS-PAGE, and processed for Western blotting using anti-DegP and anti-σ^70^ antibodies. The band corresponding to DegP is shown (Mol. wt. ∼48 kDa). Δ*degP* cultured in TBK-Brij served as a control. σ^70^ served as a loading control. The blot shown is a representative of three independent replicates. DegP levels quantified from the Western blot are shown in a bar graph. For every time point, the intensities of DegP and σ^70^ were quantified using ImageJ software. The intensity of DegP was normalized using σ^70^ as the loading control. The relative fold change in DegP levels for each time point was then calculated using DegP levels in WT grown in TBK-Brij at time point T1 as a reference. Data represent the average (±S.D.) of three independent experiments. (C) The reduced form of DegP further accumulates in oleate-grown cells in the stationary phase. Strains were grown either in TBK-Brij (-Ole) or TBK-Ole (+Ole). Cells were harvested at different phases of growth corresponding to time points T3, T4, and T5 as indicated in Fig 2A, and processed as mentioned in the legend to Fig 2F, followed by probing with an anti-DegP antibody. Δ*dsbA* and Δ*degP* strains cultured in TBK-Brij served as controls. DegP_ox_ and DegP_red_ indicate oxidized and reduced forms of DegP, respectively. Ponceau S-stained counterpart of the Western blot served as a loading control. Each blot is a representative of three independent replicates. (D) Total DegP levels in oleate-grown cells increase in the stationary phase. Strains were grown either in TBK-Brij (-Ole) or TBK-Ole (+Ole). Cells were harvested at different phases of growth corresponding to time points T3, T4, and T5 (as indicated in Fig 2A), and processed for Western blotting as mentioned in the legend to Fig 6B. The band corresponding to DegP is shown (Mol. wt. ∼48 kDa). Δ*degP* cultured in TBK-Brij served as a control. σ^70^ served as a loading control. The blot shown is a representative of four independent replicates. DegP levels quantified from the Western blot, as mentioned in the legend to Fig 6B, are shown in a bar graph. The relative fold change in DegP levels for each time point was calculated using DegP levels in WT grown in TBK-Brij at time point T3 as a reference. Data represent the average (±S.D.) of four independent experiments. (E) In oleate, total DegP levels in the stationary phase are lower in a Δ*cpxR* strain. Strains were grown either in TBK-Brij (-Ole) or TBK-Ole (+Ole). Cells were harvested at time point T5 (as indicated in Fig 2A), and processed for Western blotting as mentioned in the legend to Fig 6B. The band corresponding to DegP is shown (Mol. wt. ∼48 kDa). Δ*degP* cultured in TBK-Brij served as a control. σ^70^ served as a loading control. The blot shown is a representative of three independent replicates. DegP levels quantified from the Western blot, as mentioned in the legend to Fig 6B, are shown in a bar graph. The relative fold change in DegP levels was calculated using DegP levels in WT grown in TBK-Brij as a reference. Data represent the average (±S.D.) of three independent experiments. (F) DegP_red_ levels remarkably decrease in a Δ*cpxR* strain grown in oleate. Strains were grown either in TBK-Brij (-Ole) or TBK-Ole (+Ole). Cells were harvested at time point T5 (as indicated in Fig 2A), and processed as mentioned in the legend to Fig 2F, followed by probing with an anti-DegP antibody. DegP_ox_ and DegP_red_ indicate oxidized and reduced forms of DegP, respectively. Δ*dsbA* and Δ*degP* cultured in TBK-Brij served as controls. Ponceau S-stained counterpart of the Western blot served as a loading control. The blot shown is a representative of three independent replicates. For panels B, D, and E, the p-values were calculated using the unpaired two-tailed Student’s t test (**, P<0.01; *, P<0.05; ns, P>0.05).

In the stationary phase (time points T4 and T5), in TBK-Ole-grown cells, DegP_red_ accumulation followed a similar pattern to that of the earlier growth phases; surprisingly, its levels were even higher than at T3 (Fig 6C). Because DegP is transcriptionally regulated by the Cpx pathway (33, 34), and Cpx is induced in LCFA-grown cells in the stationary phase (where ubiquinone is sufficient), we reasoned that an upregulation of DegP by Cpx helps DegP maintain its thiol state. Assessing DegP levels from time points T3 to T5, these were indeed significantly higher in TBK-Ole-grown cells at time points T4 and T5 corresponding to the stationary phase (Fig 6D). Importantly, this increase was Cpx-dependent; deleting *cpxR*, the response regulator of the Cpx pathway, decreased DegP levels in both TBK-Brij and TBK-Ole-grown cells, with a more pronounced effect in TBK-Ole (Fig 6E). Consistent with this, DegP_red_ levels were also significantly lower in a Δ*cpxR* strain (Fig 6F). Moreover, DegP levels were significantly higher in both cells stressed with alkaline pH and those overexpressing NlpE (Figs S8 and S9), reiterating that a similar mechanism contributes to DegP activation under various stress conditions.

## Discussion

The well-established structure-driven functional model of DegP activation, derived mainly from *in vitro* studies, states that the binding of misfolded substrates transitions the inactive DegP_6_ into the functional DegP_12/24_ forms via DegP_3_ and stabilizes these active forms. Briefly, engagement of substrates with the PDZ1 domain allows them to further interact with the protease domain, initiating a cascade of conformational rearrangements in DegP that extracts the inhibitory LA loop from the active site (11, 13, 35, 36). Notably, mutating Phe residues within the protease domain, including Phe63 of the LA loop, that otherwise stabilize the hexameric state, markedly enhances proteolytic activity (37). Interestingly, the conformational change at Phe63 that WT DegP exhibits upon shift to high temperatures is observed even at temperatures as low as 20°C, when its LA loop cysteines are replaced with alanines (ΔCys DegP) (12, 14). Consistent with this, ΔCys DegP exists as DegP_3_ and DegP_12/24_ and is primed to attain DegP_12/24_ form at considerably lower substrate levels than the WT protein (14). Collectively, from these studies, one can speculate that, under *in vivo* stress conditions, DegP activation may occur due to a problem in the formation of its DSB, resulting from the compromised activity of the DSB-forming machinery, or the reduction of its DSB by thiol-containing misfolded substrates that engage DegP allosterically.

Our data provide support that, during LCFA metabolism, a redox-perturbing condition, both mechanisms operate to activate DegP; however, their contribution differs across growth phases. From exponential to entry into the stationary phase, despite constant total DegP levels, DegP_red_ levels increase with increasing cell density (Figs 6A and B). Here, ubiquinone limitation compromises DsbA functioning, which can lead to DegP activation in two ways, which are not mutually exclusive: DsbA is unable to oxidize DegP, and thiol-containing misfolded proteins (other DsbA substrates) bind to DegP. In the stationary phase, increased DegP synthesis, driven by Cpx activation, is a major factor for the increase in DegP_red_ levels (Figs 6C-F). Here, DegP activation appears predominantly substrate-driven; newly synthesized DegP molecules are likely recruited and oligomerized by substrates that accumulate in the stationary phase before DsbA can oxidize their cysteine residues. Further, re-oxidation of DegP oligomers that have already accumulated from the exponential to the stationary phase, by DsbA, also seems mechanistically unlikely; DsbA would first require DegP_12/24_ to disassemble into DegP_3_, prior to DSB formation (7). We thus propose that the stationary-phase DegP_red_ constitutes a cumulative pool comprising pre-existing DegP_red_ generated during earlier growth and newly synthesized molecules that are stabilized in their active oligomeric forms due to structural constraints and substrate-mediated effects. Whether thiol-containing substrates activate DegP via the canonical mechanism, i.e., sequential binding to the PDZ1 and protease domains, or bind the protease domain directly, where they reduce its DSB, is worth investigating. Because *in vitro* the lack of DSB primes the substrate-induced activation of DegP (14), we suggest that *in vivo* DegP_red_ facilitates its robust activation even when substrate levels alone might be insufficient to trigger activation.

In a Δ*dsbA* strain, where DegP predominantly exists as DegP_red_, it exhibits proteolytic activity even at low temperatures and is required for cellular fitness upon DTT exposure (15). This correlation between the presence of DegP as DegP_red_ and the requirement of its proteolytic activity to counter redox stress is also evident from the higher levels of DegP_red_ and the greater need for the proteolytic activity of DegP in LCFA-metabolizing cells (Figs 6A,C and 1D). DegP functions both as a chaperone and a protease, and the switch between the two activities is governed by the extent of misfolding of substrates. Whereas heavily misfolded substrates tend to get degraded by DegP, its chaperone activity suffices under milder stress (5, 11, 13). Notably, an earlier study reported that whereas overexpression of DegP_S210A_ (a DegP mutant that lacks proteolytic activity but retains its chaperone/chaperone-like activity) could support survival of Δ*degP* cells under heat shock, it failed to do so for the Δ*dsbA*Δ*degP* strain, where protein damage is extensive (38). Corroborating this, we observed that the overexpression of DegP_S210A_ supported the growth of the Δ*degP* strain in basal medium even when exposed to 7 mM DTT; however, it could support its growth only up to 6 mM DTT when cultured in LCFA-supplemented medium (Fig 1D).

DegP is widely known to relieve different kinds of stresses primarily by degrading aberrant substrates (6, 22, 24, 26); however, only under alkaline pH, it has also been shown to participate in Cpx activation (23) (Fig 5A). Here, we demonstrate that DegP also contributes to Cpx activation in cells exposed to other stress conditions where it considerably accumulates as DegP_red,_ i.e., LCFA metabolism, DTT exposure, *dsbA* deletion, and NlpE overexpression (Figs 2B-F and 6A,C) (15, 16, 31). Probing the role of DegP_red_ in detail in the LCFA condition, our data suggest that only during ubiquinone limitation, i.e., towards entry into the stationary phase, DegP_red_ functions to trigger the Cpx response; ∼40% Cpx activation that happens due to ubiquinone limitation is solely mediated via DegP (Fig 4A). Performing a detailed comparison of the mode of DegP activation and Cpx induction during alkaline stress with LCFA metabolism, our data suggest that during alkaline stress: (i) similar to LCFA-grown cells, the proteolytic activity of DegP, i.e., DegP_red_ induces Cpx response (Figs 2G and 5B), (ii) similar to stationary phase LCFA-grown cells, misfolded proteins allosterically convert the *de novo* synthesized DegP (upregulated by Cpx) into DegP_red_ (Figs 6C-F and S8), and (iii) unlike LCFA-grown cells, both DegP activation and Cpx induction occur completely in a ubiquinone-independent manner (Figs 4A,B, 5D,E and S7). Finally, because Cpx is activated upon DTT exposure, *dsbA* deletion, and NlpE overexpression (Fig 2C-E), the allosteric mode of DegP activation likely operates under these conditions as well. Figure 7 summarizes the modes of redox-mediated DegP activation under various stress conditions.

**Figure 7.**
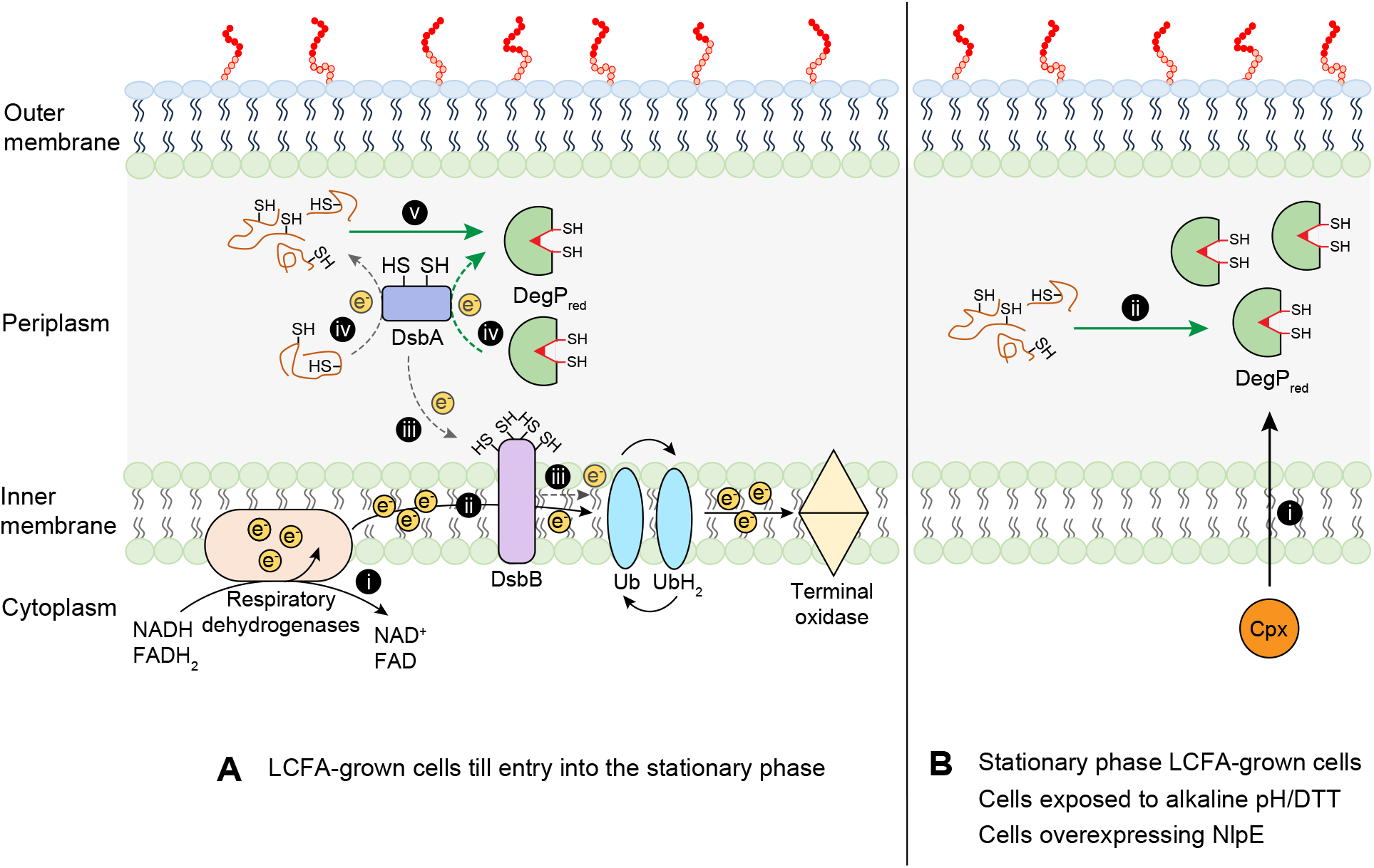
Model illustrating the dual mode of redox-mediated DegP activation under stress conditions. (A) LCFA-grown cells till entry into the stationary phase: (i) LCFA metabolism generates a large amount of NADH and FADH_2_ (ii) that increases electron flow towards the ETC (iii) rendering ubiquinone limiting to re-oxidize the DSB-forming machinery (iv) leading to the accumulation of its substrates, including DegP, in their thiol form. (v) DsbA substrates in their thiol form can also allosterically convert DegP into DegP_red_. (B) Stationary phase LCFA-grown cells, and cells exposed to alkaline pH, DTT, or overexpressing NlpE: (i) Cpx activation upregulates DegP, (ii) which is allosterically converted into DegP_red_ by thiol-containing proteins that accumulate under these stressors. The dashed arrow indicates decreased electron flow at these steps. Abbreviations: Ub, ubiquinone; UbH_2_, ubiquinol

Although under the various stress conditions tested here, DegP considerably accumulates as DegP_red_ and is involved in Cpx activation, the molecular players that activate DegP and are acted upon by the protease to induce the ESR are likely distinct across conditions. NlpE is a well-recognised inducer of the Cpx pathway and is also the signal that activates Cpx in a Δ*dsbA* strain (28-30). The involvement of DegP in activating Cpx both in NlpE overexpressing cells and in a Δ*dsbA* strain (Fig 2C and E), suggests that NlpE likely brings DegP into its thiol form and/or is acted upon by the protease to induce Cpx. In cells exposed to alkaline pH, DegP-mediated CpxP proteolysis triggers the Cpx response (23) (Fig 5B). In LCFA-metabolizing cells, although Cpx activation is partly DegP-dependent (Fig 2B), it is both NlpE- and CpxP-independent (18) (Fig 3). Thus, an as-yet-unidentified substrate/target of DegP contributes to Cpx activation in LCFA-grown cells, and because CpxP is not involved here, it likely induces Cpx via interaction with CpxA. Collectively, the multiplicity of the activators/targets of DegP enables it to integrate diverse environmental signals. Notably, around 300 secreted proteins in *E. coli* are predicted to be disulfide-bonded (39), further strengthening the idea that the redox-mediated activation of DegP positions it as a versatile, broad-spectrum protease responsive to a range of stress cues.

DegP is structurally conserved across the *Enterobacteriaceae* family of Gram-negative bacteria, particularly the LA loop architecture and the positioning of its DSB, and its proteolytic activity is a key virulence factor in several pathogens (14, 40-42). Because enteric pathogens face redox stress and metabolic shifts during infection and antibiotic exposure (43-47), a thorough understanding of redox-regulated DegP activation can help envision novel anti-virulence strategies.

## Materials and methods

### Strain, plasmids, and primers

The strains and plasmids, and primers used in this study, are listed in Tables S1 and S2, respectively. *E. coli* MG1655 was used as the WT strain for all assays. Deletion strains obtained from the Keio collection (48) were freshly transduced in the desired backgrounds using P1 phage and verified by colony PCR before final assays, to rule out genetic errors. Wherever required, the antibiotic resistance cassette was flipped out from the deletion strains using pCP20. The derivatives of Δ*cpxR*, Δ*cpxP*, Δ*degP*, Δ*cpxP*Δ*degP*, and Δ*dsbA*Δ*degP* strains were unstable as glycerol stocks and thus were always freshly made before use. The promoter-less plasmid pACYC177 was used for complementation experiments. The IPTG-inducible plasmid pRC10 was used to clone and express C-terminally sequential peptide affinity (SPA)-tagged CpxP (*cpxP*-SPA). The strain DH5α was used for cloning in plasmids pACYC177 and pRC10.

For complementation experiments, *degP*_WT_ and *degP*_S210A_ were cloned into pACYC177. Briefly, for *degP*_WT_, the coding region of *degP*, along with its promoter (325 bp upstream of the *degP* start codon), was PCR amplified and cloned into the *Bam*HI and *Hind*III sites to generate plasmid pAN01. The plasmid pDR09 harboring *the degP*_S210A_ mutant was generated by overlap extension PCR using pAN01 as the template. Plasmid pAN02 was constructed to express C-terminally SPA-tagged CpxP from the IPTG-inducible promoter of pRC10. For this, the chromosomal copy of *cpxP* was first tagged at its C-terminus with the SPA tag. Briefly, the sequence of SPA, along with the kanamycin cassette, was PCR amplified from a SPA-tagged strain obtained from the Keio collection using primers that carried sequences homologous to *cpxP*. The amplified fragment was transformed into BW25113 expressing λ Red-recombinase enzymes from pKD46 (49, 50). The recombinants were selected using kanamycin and verified by colony PCR. The SPA-tagged locus was P1 transduced into clean BW25113. The freshly transduced strain was used for amplification of *cpxP*-SPA. The amplified fragment was cloned into the *Eco*RI and *Bam*HI sites of pRC10 to generate pAN02.

### Media composition and growth conditions

The media used had the following composition: tryptone broth K (TBK)-10 g/liter Bacto tryptone, and 5 g/liter KCl, and lysogeny broth (LB)-5 g/liter Bacto yeast extract, 10 g/liter Bacto tryptone, and 5 g/liter NaCl. 100 mM potassium phosphate buffer was used to buffer TBK media at pH 7.0. Where required, media were supplemented with sodium salt of oleate (final concentration, 5 mM). A 50 mM stock of oleate was prepared in 5.0% Brij-58. Sodium oleate and Brij-58 were obtained from Sigma. Media were solidified using 1.5% (w/v) Difco agar. When required, the antibiotics, ampicillin (100 μg/ml), kanamycin (30 μg/ml), and spectinomycin (50 μg/ml), and DTT were added. For experiments performed in LB of different pH, the medium was buffered at pH 5.8, 6.8, and 8.2 by adding sodium phosphate buffer (final concentration, 100 mM) of the appropriate pH, following the protocol described in (51), with slight modifications.

Primary cultures were grown overnight (14-16 hours) in 3 ml LB. Secondary cultures were set up by re-inoculating primary cultures to an initial OD_600_ ∼0.01 either in TBK medium with or without the desired carbon source or in LB medium and grown for defined time periods. For ubiquinone supplementation experiments, 20 µM ubiquinone-8 (Avanti Polar Lipids) was added to the medium. A 20 mM stock of ubiquinone-8 was prepared in 100% ethanol (Merck). Cultures were grown in a water bath shaker at 37°C and 220 rpm. The Δ*dsbA*Δ*degP* strain was cultured at 30°C.

### Growth curve in shake flasks

Primary cultures grown overnight in LB were pelleted, washed, and resuspended in the medium used for secondary cultures. Cells were re-inoculated in 15 ml of the desired growth medium (in 125 ml flasks) to an initial OD_600_ of ∼0.01. OD_600_ was measured at defined time intervals.

### Dilution spotting

Overnight-grown cultures were pelleted, washed, and resuspended in M9 minimal medium. OD_450_ of the cultures was measured and normalized to the culture with the lowest OD_450_. Serially diluted cultures were spotted on TBK-Brij and TBK-Ole plates supplemented with different concentrations of DTT. Antibiotics were added to the plates for complementation experiments. Plates were incubated and imaged (Gel Doc GO imaging system, Bio-Rad) at different time intervals. A representative image with evident growth differences is shown in the figures.

### β-galactosidase assay

β-gal assays were performed in MG1655 *cpxP*-*lacZ* background as mentioned in (18). Briefly, cells were pelleted at different time points, washed at least four times with Z-buffer, and normalized in the same buffer to OD_450_ ∼0.5. Promoter activity was measured by monitoring β-gal expression, as described (52).

### Determination of the redox state of DegP

The *in vivo* redox state of DegP was assessed using MAL-PEG, as described previously (18). Briefly, cells were treated with trichloroacetic acid to a final concentration of 20% at the appropriate time points to prevent aerial oxidation and trap free thiols of DegP in their original state. Cultures were immediately harvested. Cells corresponding to OD_600_ ∼1.5 were used for downstream processing. Protein precipitates were pelleted by centrifugation, washed with acetone twice, dried, and resuspended in a freshly prepared buffer containing 100 mM Tris-Cl (pH 7.5), 10 mM EDTA, 1% SDS, and 2 mM MAL-PEG. Samples were subjected to 25°C incubation in the dark for 1 hour at 1,400 revolutions/min. Proteins were resolved on 10% non-reducing SDS-PAGE gels, transferred to nitrocellulose membrane, and processed for Western blotting.

### Determination of the expression levels of DegP and CpxP-SPA

Cultures were harvested at the desired time points, and cells corresponding to OD_600_ ∼0.5 were pelleted at 6,000 rpm for 5 min. Proteins were resolved on 10% SDS-PAGE gels for DegP and 15% SDS-PAGE gels for CpxP-SPA, transferred to nitrocellulose membrane, and processed for Western blotting.

### Western blotting

The expression levels and the redox state of proteins were determined by Western blot analysis. Samples were run on SDS-PAGE gels and transferred to nitrocellulose membrane, followed by membrane blocking with 5% (w/v) skimmed milk overnight at 4°C and probing with anti-DegP (1:10,000) or anti-FLAG (1:3,300, Sigma) primary antibody and HRP-conjugated anti-rabbit (1:5,000, Sigma) for DegP or HRP-conjugated anti-mouse (1:5,000, Sigma) secondary antibody for CpxP-SPA. Blots were developed using the SuperSignal West Dura Extended Duration Substrate (Pierce), and the chemiluminescence signals were either imaged using X-ray films or were detected by ImageQuant LAS 4000. The bands in the Western blot were quantified using ImageJ 1.54D (53), as required.

## Data Availability

All data are available in the main article and its supporting information.

## Acknowledgments

We thank Jean-François Collet for plasmids and Manjula Reddy for anti-DegP antibody. We thank members of the Chaba lab and Jogender Singh for discussions and critical reading of the manuscript. This work was supported by the Department of Biotechnology (DBT)/Wellcome Trust India Alliance senior fellowship/grant IA/S/21/2/505907 and Ministry of Education-Scheme for Transformational and Advanced Research in Sciences (STARS) grant MoE-STARS/STARS-1/296, and partially supported by the Department of Science and Technology-Science and Engineering Research Board (DST-SERB) grant CRG/2018/000833, DBT grant BT/PR34553/BRB/10/1846/2020, and funds from IISER Mohali (to R.C.). D.R. acknowledges IISER Mohali for fellowship support for doctoral work.

## Notes

### Competing Interest Statement

The authors have declared no competing interest.

## References

1. T. J. Silhavy, D. Kahne, S. Walker, The bacterial cell envelope. Cold Spring Harbor perspectives in biology 2, a000414 (2010).

2. J. Skorko-Glonek et al., HtrA heat shock protease interacts with phospholipid membranes and undergoes conformational changes. The Journal of biological chemistry 272, 8974–8982 (1997).

3. T. Clausen, M. Kaiser, R. Huber, M. Ehrmann, HTRA proteases: regulated proteolysis in protein quality control. Nature reviews. Molecular cell biology 12, 152–162 (2011).

4. T. Clausen, C. Southan, M. Ehrmann, The HtrA family of proteases: implications for protein composition and cell fate. Molecular cell 10, 443–455 (2002).

5. C. Spiess, A. Beil, M. Ehrmann, A temperature-dependent switch from chaperone to protease in a widely conserved heat shock protein. Cell 97, 339–347 (1999).

6. X. Fu et al., DegP functions as a critical protease for bacterial acid resistance. The FEBS journal 285, 3525–3538 (2018).

7. J. Jiang et al., Activation of DegP chaperone-protease via formation of large cage-like oligomers upon binding to substrate proteins. Proceedings of the National Academy of Sciences of the United States of America 105, 11939–11944 (2008).

8. M. Merdanovic et al., Determinants of structural and functional plasticity of a widely conserved protease chaperone complex. Nature structural & molecular biology 17, 837–843 (2010).

9. S. Kim, R. T. Sauer, Cage assembly of DegP protease is not required for substrate-dependent regulation of proteolytic activity or high-temperature cell survival. Proceedings of the National Academy of Sciences of the United States of America 109, 7263–7268 (2012).

10. T. Krojer, M. Garrido-Franco, R. Huber, M. Ehrmann, T. Clausen, Crystal structure of DegP (HtrA) reveals a new protease-chaperone machine. Nature 416, 455–459 (2002).

11. T. Krojer et al., Structural basis for the regulated protease and chaperone function of DegP. Nature 453, 885–890 (2008).

12. A. Sobiecka-Szkatula et al., Temperature-induced conformational changes within the regulatory loops L1-L2-LA of the HtrA heat-shock protease from Escherichia coli. Biochimica et biophysica acta 1794, 1573–1582 (2009).

13. J. Ortega, J. Iwanczyk, A. Jomaa, Escherichia coli DegP: a structure-driven functional model. Journal of bacteriology 191, 4705–4713 (2009).

14. T. Koper et al., Analysis of the link between the redox state and enzymatic activity of the HtrA (DegP) protein from Escherichia coli. PloS one 10, e0117413 (2015).

15. J. Skorko-Glonek, A. Sobiecka-Szkatula, J. Narkiewicz, B. Lipinska, The proteolytic activity of the HtrA (DegP) protein from Escherichia coli at low temperatures. Microbiology (Reading) 154, 3649–3658 (2008).

16. J. Skorko-Glonek, A. Sobiecka-Szkatula, B. Lipinska, Characterization of disulfide exchange between DsbA and HtrA proteins from Escherichia coli. Acta biochimica Polonica 53, 585–589 (2006).

17. B. Manta, D. Boyd, M. Berkmen, Disulfide Bond Formation in the Periplasm of Escherichia coli. EcoSal Plus 8 (2019).

18. K. Jaswal, M. Shrivastava, D. Roy, S. Agrawal, R. Chaba, Metabolism of long-chain fatty acids affects disulfide bond formation in Escherichia coli and activates envelope stress response pathways as a combat strategy. PLoS genetics 16, e1009081 (2020).

19. M. Shrivastava et al., An envelope stress response governs long-chain fatty acid metabolism via a small RNA to maintain redox homeostasis in Escherichia coli. bioRxiv : the preprint server for biology 10.1101/2024.10.18.618624, 2024.2010.2018.618624 (2024).

20. K. Jaswal, M. Shrivastava, R. Chaba, Revisiting long-chain fatty acid metabolism in Escherichia coli: integration with stress responses. Current genetics 67, 573–582 (2021).

21. M. Shrivastava, D. Roy, R. Chaba, Long-chain fatty acids as nutrients for Gram-negative bacteria: stress, proliferation, and virulence. Current opinion in microbiology 85, 102609 (2025).

22. S. Zhang et al., Degp degrades a wide range of substrate proteins in Escherichia coli under stress conditions. The Biochemical journal 476, 3549–3564 (2019).

23. D. R. Buelow, T. L. Raivio, Cpx signal transduction is influenced by a conserved Nterminal domain in the novel inhibitor CpxP and the periplasmic protease DegP. Journal of bacteriology 187, 6622–6630 (2005).

24. X. Ge et al., DegP primarily functions as a protease for the biogenesis of beta-barrel outer membrane proteins in the Gram-negative bacterium Escherichia coli. The FEBS journal 281, 1226–1240 (2014).

25. J. Skorko-Glonek et al., The Escherichia coli heat shock protease HtrA participates in defense against oxidative stress. Molecular & general genetics : MGG 262, 342–350 (1999).

26. D. D. Isaac, J. S. Pinkner, S. J. Hultgren, T. J. Silhavy, The extracytoplasmic adaptor protein CpxP is degraded with substrate by DegP. Proceedings of the National Academy of Sciences of the United States of America 102, 17775–17779 (2005).

27. P. N. Danese, T. J. Silhavy, CpxP, a stress-combative member of the Cpx regulon. Journal of bacteriology 180, 831–839 (1998).

28. A. Delhaye, G. Laloux, J. F. Collet, The Lipoprotein NlpE Is a Cpx Sensor That Serves as a Sentinel for Protein Sorting and Folding Defects in the Escherichia coli Envelope. Journal of bacteriology 201 (2019).

29. J. Marotta, K. L. May, C. Y. Bae, M. Grabowicz, Molecular insights into Escherichia coli Cpx envelope stress response activation by the sensor lipoprotein NlpE. Molecular microbiology 119, 586–598 (2023).

30. K. L. May, K. M. Lehman, A. M. Mitchell, M. Grabowicz, A Stress Response Monitoring Lipoprotein Trafficking to the Outer Membrane. mBio 10 (2019).

31. J. Skorko-Glonek et al., The N-terminal region of HtrA heat shock protease from Escherichia coli is essential for stabilization of HtrA primary structure and maintaining of its oligomeric structure. Biochimica et biophysica acta 1649, 171–182 (2003).

32. A. M. Mitchell, T. J. Silhavy, Envelope stress responses: balancing damage repair and toxicity. Nature reviews. Microbiology 17, 417–428 (2019).

33. T. L. Raivio, Everything old is new again: an update on current research on the Cpx envelope stress response. Biochimica et biophysica acta 1843, 1529–1541 (2014).

34. J. Pogliano, A. S. Lynch, D. Belin, E. C. Lin, J. Beckwith, Regulation of Escherichia coli cell envelope proteins involved in protein folding and degradation by the Cpx two-component system. Genes & development 11, 1169–1182 (1997).

35. S. Kim, R. A. Grant, R. T. Sauer, Covalent linkage of distinct substrate degrons controls assembly and disassembly of DegP proteolytic cages. Cell 145, 67–78 (2011).

36. T. Krojer et al., Interplay of PDZ and protease domain of DegP ensures efficient elimination of misfolded proteins. Proceedings of the National Academy of Sciences of the United States of America 105, 7702–7707 (2008).

37. D. Figaj et al., The LA loop as an important regulatory element of the HtrA (DegP) protease from Escherichia coli: structural and functional studies. The Journal of biological chemistry 289, 15880–15893 (2014).

38. J. Skorko-Glonek, E. Laskowska, A. Sobiecka-Szkatula, B. Lipinska, Characterization of the chaperone-like activity of HtrA (DegP) protein from Escherichia coli under the conditions of heat shock. Archives of biochemistry and biophysics 464, 80–89 (2007).

39. R. J. Dutton, D. Boyd, M. Berkmen, J. Beckwith, Bacterial species exhibit diversity in their mechanisms and capacity for protein disulfide bond formation. Proceedings of the National Academy of Sciences of the United States of America 105, 11933–11938 (2008).

40. P. Redford, R. A. Welch, Role of sigma E-regulated genes in Escherichia coli uropathogenesis. Infection and immunity 74, 4030–4038 (2006).

41. C. Lewis et al., Salmonella enterica Serovar Typhimurium HtrA: regulation of expression and role of the chaperone and protease activities during infection. Microbiology (Reading) 155, 873–881 (2009).

42. J. Skorko-Glonek et al., HtrA protease family as therapeutic targets. Current pharmaceutical design 19, 977–1009 (2013).

43. S. Sultana et al., Redox-Mediated Inactivation of the Transcriptional Repressor RcrR is Responsible for Uropathogenic Escherichia coli’s Increased Resistance to Reactive Chlorine Species. mBio 13, e0192622 (2022).

44. A. V. Nair, A. Singh, R. S. Rajmani, D. Chakravortty, Salmonella Typhimurium employs spermidine to exert protection against ROS-mediated cytotoxicity and rewires host polyamine metabolism to ameliorate its survival in macrophages. Redox biology 72, 103151 (2024).

45. A. Vazquez-Torres, F. C. Fang, Salmonella evasion of the NADPH phagocyte oxidase. Microbes and infection 3, 1313–1320 (2001).

46. M. A. Lobritz et al., Antibiotic efficacy is linked to bacterial cellular respiration. Proceedings of the National Academy of Sciences of the United States of America 112, 8173–8180 (2015).

47. M. Lang, A. Carvalho, Z. Baharoglu, D. Mazel, Aminoglycoside uptake, stress, and potentiation in Gram-negative bacteria: new therapies with old molecules. Microbiology and molecular biology reviews : MMBR 87, e0003622 (2023).

48. T. Baba et al., Construction of Escherichia coli K-12 in-frame, single-gene knockout mutants: the Keio collection. Molecular systems biology 2, 2006 0008 (2006).

49. K. A. Datsenko, B. L. Wanner, One-step inactivation of chromosomal genes in Escherichia coli K-12 using PCR products. Proceedings of the National Academy of Sciences of the United States of America 97, 6640–6645 (2000).

50. S. Datta, N. Costantino, D. L. Court, A set of recombineering plasmids for gram-negative bacteria. Gene 379, 109–115 (2006).

51. J. Sambrook, Molecular cloning : a laboratory manual (Third edition. Cold Spring Harbor, N.Y. : Cold Spring Harbor Laboratory Press, [2001] ©2001, 2001).

52. J. H. Miller, Experiments in Molecular Genetics. Cold Spring Harbor Laboratory, Cold Spring Harbor, New York. (1972).

53. C. A. Schneider, W. S. Rasband, K. W. Eliceiri, NIH Image to ImageJ: 25 years of image analysis. Nature methods 9, 671–675 (2012).

54. R. Malpica, B. Franco, C. Rodriguez, O. Kwon, D. Georgellis, Identification of a quinone-sensitive redox switch in the ArcB sensor kinase. Proceedings of the National Academy of Sciences of the United States of America 101, 13318–13323 (2004).

