## Supporting information for "Distinct modes of redox-mediated DegP activation govern its role as a general protease"

**This PDF file includes:**

Figures S1 to S9
Tables S1 and S2

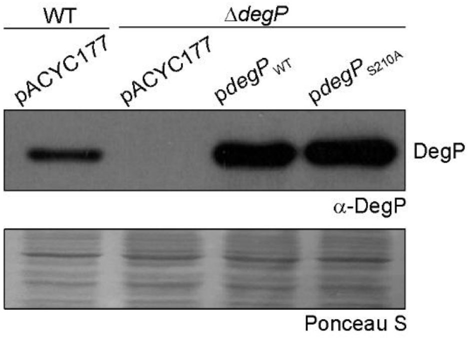


**Fig. S1.** *$DegP_{WT}$ and $DegP_{S210A}$ exhibit similar expression levels.* WT and Δ*degP* strains transformed with either pACYC177, p*degP*$_{WT}$, or p*degP*$_{S210A}$ were grown in TBK-Ole, and cultures were harvested. The band corresponding to DegP (~48 kDa) is shown. Δ*degP* transformed with pACYC177 served as a control. Ponceau S-stained counterpart of the Western blot served as a loading control. The blot is a representative of two independent replicates.

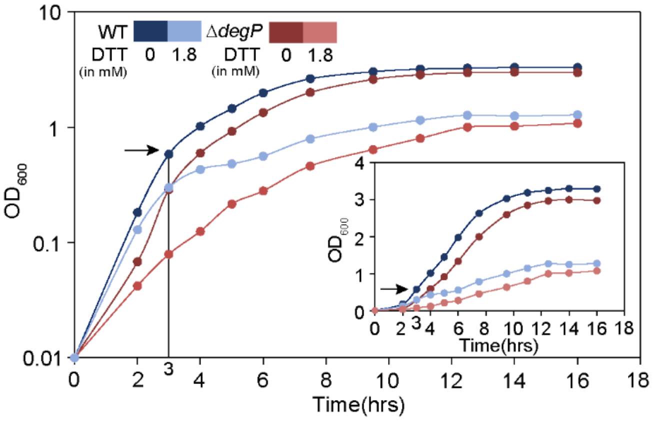


**Fig. S2.** *Growth curve of WT and ΔdegP strains in TBK with or without DTT.* Strains were grown in TBK medium with or without supplementation of 1.8 mM DTT. $OD_{600}$ of the cultures was measured, and growth curves were plotted on a semi-logarithmic scale. The experiment was done twice. A representative dataset is shown. The arrow indicates the time point where cultures were harvested to check the effect of *degP* deletion on Cpx activation in Fig 2D. *Inset*: The above growth curves were also plotted on a linear scale.

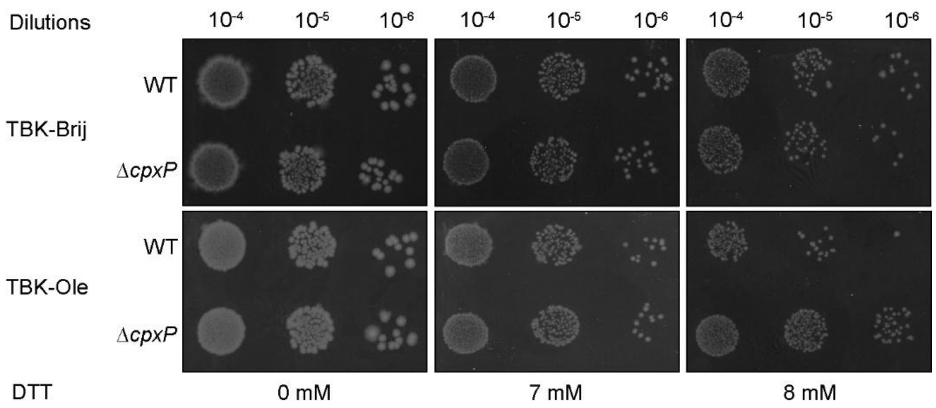


**Fig. S3.** *Deletion of cpxP confers a growth advantage to oleate-grown cells exposed to thiol stress.* Strains were spotted either on solid TBK-Brij or TBK-Ole media with or without the indicated concentrations of DTT. The experiment was performed three times. A representative dataset is shown.

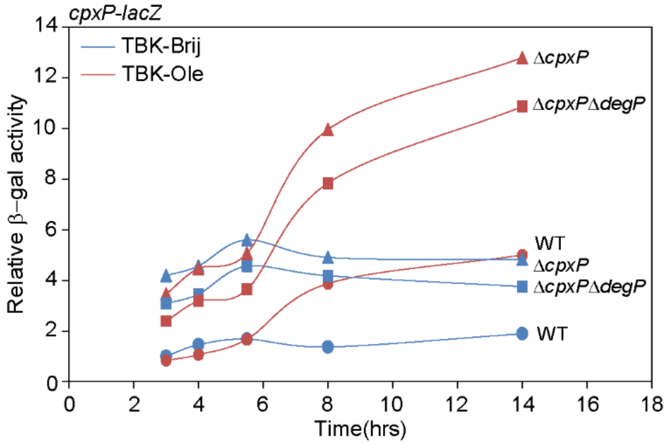


**Fig. S4.** *In oleate-grown cells, DegP-mediated activation of Cpx is CpxP-independent.* Strains carrying *cpxP-lacZ* transcriptional reporter were grown either in TBK-Brij or TBK-Ole. Cultures were harvested at different phases of growth corresponding to the time points as indicated in Fig 2A, and β-gal activity was measured. Data were normalized to the β-gal activity of WT grown in TBK-Brij at time point T1. The β-gal activity (in Miller units) of WT *cpxP-lacZ* in TBK-Brij at time point T1 was 45. Two independent experiments were performed; although the fold change varied across biological replicates, the trend observed was the same. Data from another independent experiment is shown in Fig 3B.

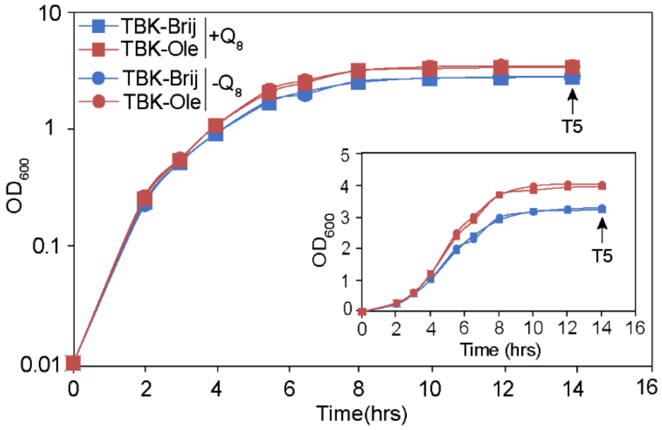


**Fig. S5.** *Growth curve of WT in TBK-Brij and TBK-Ole with or without ubiquinone-8 supplementation.* WT was grown in TBK-Brij and TBK-Ole supplemented either with 20 μM ubiquinone-8 ($Q_8$+) or 0.1% ethanol ($Q_8$-). $OD_{600}$ of the cultures was measured, and growth curves were plotted on a semi-logarithmic scale. The experiment was done twice. A representative dataset is shown. The arrow indicates time point T5 where cultures were harvested for various experiments (Fig 4A and B). *Inset:* The above growth curves were also plotted on a linear scale.

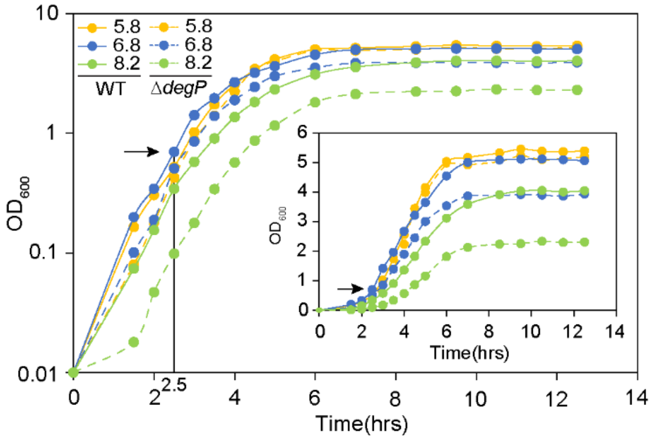


**Fig. S6.** *Growth curve of WT in LB buffered at different pH.* WT was grown in LB medium buffered at pH 5.8, 6.8, and 8.2. $OD_{600}$ of the cultures was measured, and growth curves were plotted on a semi-logarithmic scale. The experiment was done twice. A representative dataset is shown. The arrow indicates the exponential phase where cultures were harvested for various experiments (Figs 5 and S7). *Inset:* The above growth curves were also plotted on a linear scale.

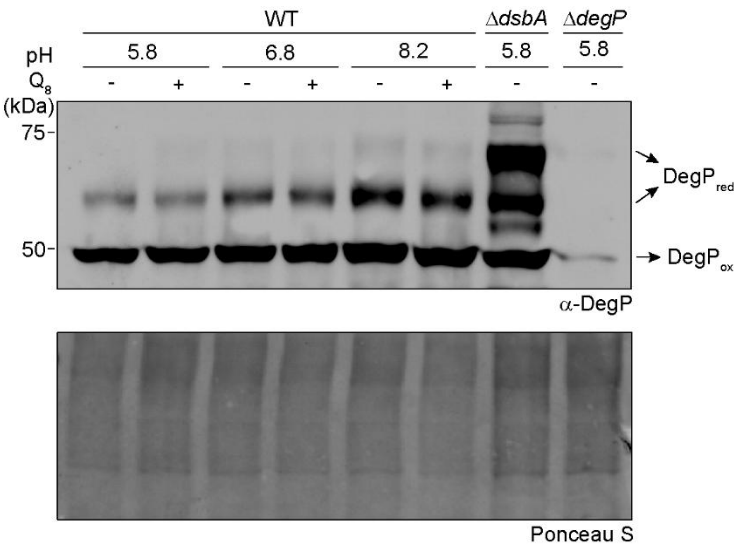


**Fig. S7.** *DegP$_{red}$ levels do not change in cells grown in alkaline pH upon ubiquinone supplementation.* Strains were grown in LB of different pH, as indicated. The media contained either 20 μM ubiquinone-8 ($Q_8$+) or 0.1% ethanol ($Q_8$-) as mentioned. Cells were harvested in the exponential phase (as indicated in Fig S6), and processed as mentioned in the legend to Fig 2F, followed by probing with an anti-DegP antibody. Δ*dsbA* and Δ*degP* strains cultured in LB of pH 5.8 served as controls. DegP$_{ox}$ and DegP$_{red}$ indicate oxidized and reduced forms of DegP, respectively. Ponceau S-stained counterpart of the Western blot served as a loading control.

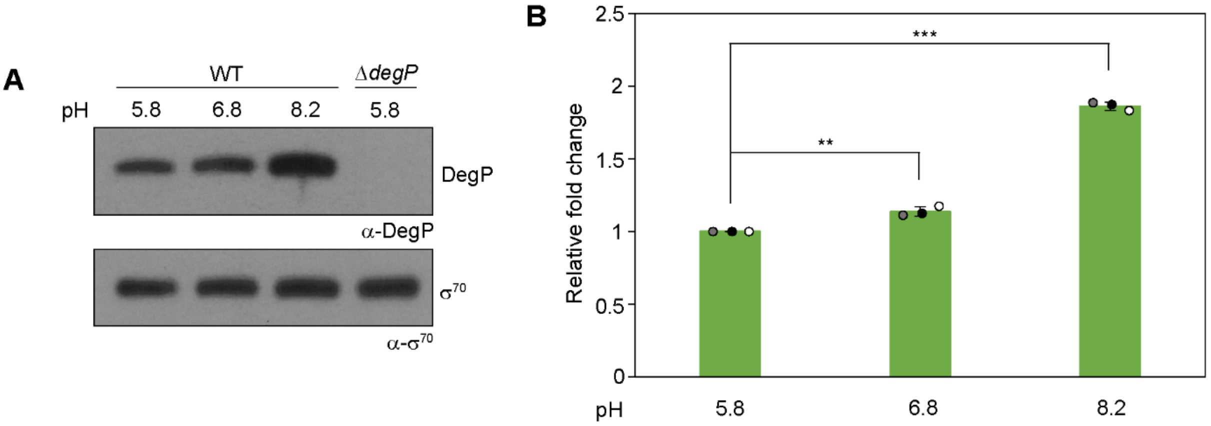


**Fig. S8.** *DegP levels gradually increase in cells exposed to increasing pH.* WT was grown in LB medium buffered at pH 5.8, 6.8, and 8.2. Cells were harvested in the exponential phase (as indicated in Fig S6), and processed for Western blotting as mentioned in the legend to Fig 6B. The band corresponding to DegP is shown (Mol. wt. ~48 kDa). Δ*degP* cultured in pH 5.8 served as a control. $\sigma^{70}$ served as a loading control. The blot shown is a representative of three independent replicates (A). DegP levels were quantified from the Western blot as mentioned in the legend to Fig 6B. The relative fold change in DegP levels for each pH was calculated using DegP levels in WT grown in pH 5.8 as a reference. Data represent the average (±S.D.) of three independent experiments. The p-values were calculated using the unpaired two-tailed Student's t-test (***, P<0.001; **, P<0.01) (B).

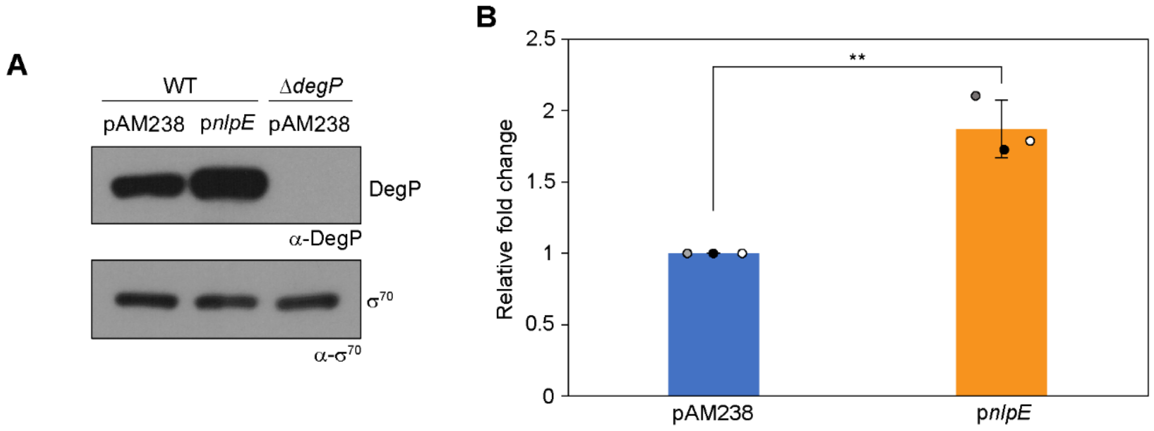


**Fig. S9.** *DegP levels significantly increase in cells overexpressing NlpE.* Strains were transformed with either pAM238 or p*nlpE* and were grown in LB medium supplemented with 0.1 mM IPTG. Cultures were harvested in the exponential phase, and processed for Western blotting as mentioned in the legend to Fig 6B. The band corresponding to DegP is shown (Mol. wt. ~48 kDa). Δ*degP* transformed with pAM238 served as a control. $\sigma^{70}$ served as a loading control. The blot shown is a representative of three independent replicates (A). DegP levels were quantified from the Western blot as mentioned in the legend to Fig 6B. The relative fold change in DegP levels in WT transformed with p*nlpE* was calculated using DegP levels in WT transformed with pAM238 as a reference. Data represent the average (±S.D.) of three independent experiments. The p-values were calculated using the unpaired two-tailed Student’s t-test (**, $P<0.01$) (B).

**Table S1**: Strains and plasmids used in this study

| Strains/Plasmids | Relevant genotype | Source (reference) |
|---|---|---|
| **Strains** | | |
| DH5α | F- Δ(*argF-lac*)*169* Φ80d*lacZ58*(M15) *glnX44*(AS) $\lambda^-$ *rfbC1 gyrA96* (Nal$^r$) *recA1 endA1 spoT1 thiE1 hsdR17* | New England Biolabs |
| BW25113 | F- Δ(*araD-araB*)*567* Δ*lacZ4787*(::rrnB-3) $\lambda^-$ *rph-1* Δ(*rhaD-rhaB*)*568 hsdR514* | *E. coli* Genetic Stock Centre |
| MG1655 | $F^-$ $\lambda^-$ *rph-1* | *E. coli* Genetic Stock Centre |
| SEA4166 | MG1655 Δ*lacX74* λ*RS88* [P*cpxP-lacZ*] | Ades Lab |
| BW25113 Δ*degP* | BW25113 *degP*::*kan*, Kan$^r$ | Keio collection (1) |
| BW25113 Δ*dsbA* | BW25113 *dsbA*::*kan*, Kan$^r$ | Keio collection (1) |
| BW25113 Δ*cpxP* | BW25113 *cpxP*::*kan*, Kan$^r$ | Keio collection (1) |
| BW25113 Δ*cpxR* | BW25113 *cpxR*::*kan*, Kan$^r$ | Keio collection (1) |
| Freshly made | P1 (BW25113 *degP*::*kan*) X MG1655, Kan$^r$ | This work |
| RC18012 | P1 (BW25113 *dsbA*::*kan*) X MG1655, Kan$^r$ | This work |
| Freshly made | P1 (BW25113 *cpxP*::*kan*) X MG1655, Kan$^r$ | This work |
| Freshly made | P1 (BW25113 *cpxR*::*kan*) X MG1655, Kan$^r$ | This work |
| Freshly made | P1 (BW25113 *degP*::*kan*) X SEA4166, Kan$^r$ | This work |
| RC18156 | P1 (BW25113 *dsbA*::*kan*) X SEA4166, Kan$^r$ | This work |
| Freshly made | P1 (BW25113 *cpxP*::*kan*) X SEA4166, Kan$^r$ | This work |
| RC18157 | SEA4166 Δ*dsbA* (*kan* cassette flipped from RC18156) | This work |
| Freshly made | P1 (BW25113 *degP*::*kan*) X RC18157, Kan$^r$ | This work |
| Freshly made | SEA4166 Δ*degP* (*kan* cassette flipped from SEA4166 *degP*::*Kan*) | This work |
| Freshly made | P1 (BW25113 *cpxP*::*kan*) X SEA4166 Δ*degP*, Kan$^r$ | This work |
| **Plasmids** | | |
| pAM238 | pSC101 *ori*, inducible $P_{lac}$, Spec$^r$ | Collet Lab (2) |
| pAM238-*nlpE* (p*nlpE*) | pSC101 *ori*, $P_{lac}$-*nlpE*, Spec$^r$ | Collet Lab (2) |
| pACYC177 | p15A *ori*, Amp$^r$ Kan$^r$ | New England Biolabs |
| pAN01 | *degP* promoter and $degP_{WT}$ in pACYC177, Amp$^r$ | This work |
| pDR09 | *degP* promoter and $degP_{S210A}$ in pACYC177, Amp$^r$ | This work |
| pKD46 | pSC101 *ori araC repA101*(Ts) $P_{araBAD}$-λred, Amp$^r$ | (3) |
| pRC10 | pBR322 *ori*, -10 box of $P_{trc}$ changed to $P_{lac}$ in pTrc99a, Δ*Nco*I, Amp$^r$ | (4) |
| pAN02 | *cpxP*-SPA in pRC10, Amp$^r$ | This work |
| pCP20 | pSC101 *ori cI857* λ-$P_R$ *flp* ts Amp$^r$ Cam$^r$ | (3) |

**Table S2**: Primers used in this study

| Primers | Sequence (5' to 3') | Purpose | Source (reference) |
|---|---|---|---|
| AN09 | ACCGGGATCCGCGCTTATTCCACAAACTCTCGA | Forward primer for cloning *degP* in pACYC177 | This work |
| AN10 | CGTAAGCTTGCACGGCTTAGCATAAGGAAGTAC | Reverse primer for cloning *degP* in pACYC177 | This work |
| AN19 | ATCAACCGTGGTAACGCAGGTGGTGCGCTGGTT | Internal mutagenic primer for creating the S210A mutation in *degP* | This work |
| AN20 | AACCAGCGCACCACCTGCGTTACCACGGTTGAT | Internal mutagenic primer for creating the S210A mutation in *degP* | This work |
| AN11 | GAGCTGATGAGCAATTTCCGTTG | Sequencing/verification primer for cloning in pACYC177 | This work |
| AN12 | AATATTGTTGATGCGCTGGCAGTG | Sequencing/verification primer for cloning in pACYC177 | This work |
| AN05 | GCAAAAAAGTTCATCGTTGAAGCTATTGAGTAGTAGCAACTCACGTTCCCAGTCCATGGAAAAGAGAAGATGG | Forward primer for tagging *cpxP* on the chromosome with SPA | This work |
| AN06 | CATGTGGGGGAAGACAGGGATGGTGTCTATGGCAAGGAAAACAGGGTTTACATATGAATATCCTCCTTAG | Reverse primer for tagging *cpxP* on the chromosome with SPA | This work |
| DR015 | ACCGGAATTCGAAGGAGATATACATATGCGCATAGTTACCGCTGCC | Forward primer for cloning *cpxP*-SPA in pRC10 | This work |
| DR016 | CGTGGATCCCTACTTGTCATCGTCATCCTTGTAGTCG | Reverse primer for cloning *cpxP*-SPA in pRC10 | This work |
| BS25 | GCTGTGGTATGGCTGTGCAGG | Sequencing/verification primer for cloning in pRC10 | (5) |
| BS26 | GCCAGGCAAATTCTGTTTTATCAG | Sequencing/verification primer for cloning in pRC10 | (5) |
| SAK1 | GAGGCTATTCGGCTATGACTG | Forward primer specific to kanamycin cassette for verification of chromosomal *cpxP*-SPA | (5) |
| SAK2 | TTCCATCCGAGTACGTGCTC | Reverse primer specific to kanamycin cassette for verification of chromosomal *cpxP*-SPA | (5) |

*Restriction sites are underlined

**SI References**


1. T. Baba *et al.*, Construction of *Escherichia coli* K-12 in-frame, single-gene knockout mutants: the Keio collection. *Molecular systems biology* **2**, 2006 0008 (2006).
2. A. Delhaye, G. Laloux, J. F. Collet, The Lipoprotein NlpE Is a Cpx Sensor That Serves as a Sentinel for Protein Sorting and Folding Defects in the *Escherichia coli* Envelope. *Journal of bacteriology* **201** (2019).
3. K. A. Datsenko, B. L. Wanner, One-step inactivation of chromosomal genes in *Escherichia coli* K-12 using PCR products. *Proceedings of the National Academy of Sciences of the United States of America* **97**, 6640-6645 (2000).
4. R. Chaba, I. L. Grigorova, J. M. Flynn, T. A. Baker, C. A. Gross, Design principles of the proteolytic cascade governing the σE-mediated envelope stress response in *Escherichia coli*: keys to graded, buffered, and rapid signal transduction. *Genes & development* **21**, 124-136 (2007).
5. B. Singh *et al.*, Molecular and Functional Insights into the Regulation of D-Galactonate Metabolism by the Transcriptional Regulator DgoR in *Escherichia coli*. *Journal of bacteriology* **201** (2019).
